# Allosteric modulation of proton binding confers Cl^-^ activation and glutamate selectivity to vesicular glutamate transporters

**DOI:** 10.1101/2024.09.06.609381

**Authors:** Bart Borghans, Daniel Kortzak, Piersilvio Longo, Jan-Philipp Machtens, Christoph Fahlke

## Abstract

Vesicular glutamate transporters (VGLUTs) fill synaptic vesicles with glutamate and remove luminal Cl^-^ via an additional anion channel mode. Both of these transport functions are stimulated by luminal acidification, luminal-positive membrane potential, and luminal Cl^-^. We studied VGLUT1 transporter/channel activation using a combination of heterologous expression, cellular electrophysiology, fast solution exchange, and mathematical modeling. Cl^-^ channel gating can be described with a kinetic scheme that includes two protonation sites and distinct opening, closing, and Cl^-^-binding rates for each protonation state. Cl^-^ binding promotes channel opening by modifying the p*K*_a_ values of the protonation sites and rates of pore opening and closure. VGLUT1 transports glutamate and aspartate at distinct stoichiometries: H^+^-glutamate exchange at 1:1 stoichiometry and aspartate uniport. Neurotransmitter transport with variable stoichiometry can be described with an alternating access model that assumes that transporters without substrate translocate in the doubly protonated state to the inward-facing conformation and return with the bound amino acid substrate as either singly or doubly protonated. Glutamate, but not aspartate, promotes the release of one proton from inward-facing VGLUT1, resulting in preferential H^+^-coupled glutamate exchange. Cl^-^ stimulates glutamate transport by making the glutamate-binding site accessible to cytoplasmic glutamate and by facilitating transitions to the inward-facing conformation after outward substrate release. We conclude that allosteric modification of transporter protonation by Cl^-^ is crucial for both VGLUT1 transport functions.

## Introduction

Glutamate is the major excitatory neurotransmitter in the mammalian central nervous system. It is released from presynaptic nerve terminals via exocytosis of synaptic vesicles that have been filled by three vesicular glutamate transporters: VGLUT1, VGLUT2, or VGLUT3 (Eriksen, Chang et al. 2016, Omote, Miyaji et al. 2016, Farsi, Jahn et al. 2017). VGLUTs employ electrochemical H^+^ gradients generated by V-type ATPases to accumulate glutamate via stoichiometrically coupled H^+^-glutamate exchange (Kolen, Borghans et al. 2023). H^+^-anion transport is partially uncoupled for aspartate, and the difference in coupling stoichiometry permits selective glutamate accumulation in synaptic vesicles (Kolen, Borghans et al. 2023). VGLUTs also function as anion channels that mediate Cl^-^ diffusion out of the synaptic vesicle. The distinct transport mechanisms for glutamate and Cl^-^ permit VGLUTs to harness the outwardly directed Cl^-^ gradient to depolarize the vesicular membrane potential and stimulate glutamate accumulation (Kolen, Borghans et al. 2023) and to exchange vesicular Cl^-^ for glutamate during synaptic vesicle filling (Schenck, Wojcik et al. 2009, Takamori 2016, Martineau, Guzman et al. 2017).

Glutamate transport rates are modified by the transmembrane voltage, acidic luminal pH, and chloride concentration ([Cl^-^]) in the lumen and cytoplasm. In synaptic vesicle preparations or in liposomes containing purified and reconstituted transporters, glutamate uptake is low in the absence of Cl^-^, but increases to maximum values at a [Cl^-^] of 4 mM. Higher concentrations inhibit glutamate uptake into synaptic vesicles, resulting in a biphasic Cl^-^ dependence (Naito and Ueda 1985, Maycox, Deckwerth et al. 1988, Moriyama and Yamamoto 1995). Experiments in proteoliposomes (Juge, Gray et al. 2010) and heterologous expression systems (Eriksen, Chang et al. 2016) have demonstrated that—at high concentrations—Cl^-^ influx via VGLUT Cl^-^ channels affects electrogenic VGLUT glutamate transport and anion channel currents by hyperpolarizing the membrane potential. However, Cl^-^ also modulates VGLUT function as an allosteric activator, as shown in experiments using endosomal patch clamp recordings in transfected cells (Chang, Eriksen et al. 2018). We combined heterologous expression of surface membrane-targeted mutant VGLUT1s and whole-cell patch clamp with fast H^+^ and Cl^-^ application and mathematical modeling to describe allosteric VGLUT activation. Our results provide the first mechanistic insights into the opening as well as the voltage and ligand gating of VGLUT anion channels and into substrate selectivity and regulation of VGLUT neurotransmitter transport.

## Results

### External Cl^-^ modifies the amplitude and kinetics of VGLUT1_PM_ Cl^-^ currents

VGLUTs require external acidification for glutamate transport and anion channel activity and are activated by luminal and cytoplasmic [Cl^-^] (Schenck, Wojcik et al. 2009). Since only luminal [Cl^-^] undergoes major modification during synaptic function (Martineau, Guzman et al. 2017, Kolen, Borghans et al. 2023), we restricted our analysis to regulation by luminal Cl^-^. We first focused on measuring Cl^-^ currents in the absence of the transport substrate glutamate. In all experiments, we expressed a surface membrane insertion-optimized mutant of VGLUT1 (VGLUT1_PM_) in HEK293T cells, followed by whole-cell patch clamping (Kolen, Borghans et al. 2023). As the luminal side of surface membrane-optimized VGLUT1_PM_ faces the external solution, the luminal [Cl^-^] can be varied by changing the external [Cl^-^], outside the cell ([Cl^-^]_o_).

Figure 1 A shows the current responses of HEK293T cells with heterologously expressed VGLUT1_PM_ to voltage steps between -160 mV and +60 mV at three external [Cl^-^] (0, 40, and 140 mM) and an external pH of 5.5. In the complete absence of external Cl^-^, currents are small but clearly above background. Increased [Cl^-^]_o_ enhances current amplitudes and accelerates the time course of activation (Figure 1 A). Figure 1 B depicts a plot of late current amplitudes versus external [Cl^-^], which is a good fit for a Michaelis–Menten relationship providing an apparent *K*_M_ of 28.3 ± 0.7 mM (mean and 95% confidence interval, with bootstrapped global fit and sampling of 1000, n = 11). Under all test conditions, VGLUT1 currents are pH dependent, with zero current amplitude at neutral external pH. Figure 1 C provides the pH dependence for two different [Cl^-^]_o_ at -160 mV, with indistinguishable p*K*_M_ values of 5.3 ± 0.003 (n = 10, Hill coefficient = 1.2) at [Cl^-^]_o_ = 0 mM and 5.4 ± 0.003 (n = 11, Hill coefficient = 1.1, p = 0.082) at [Cl^-^]_o_ = 140 mM.

**Figure 1:**
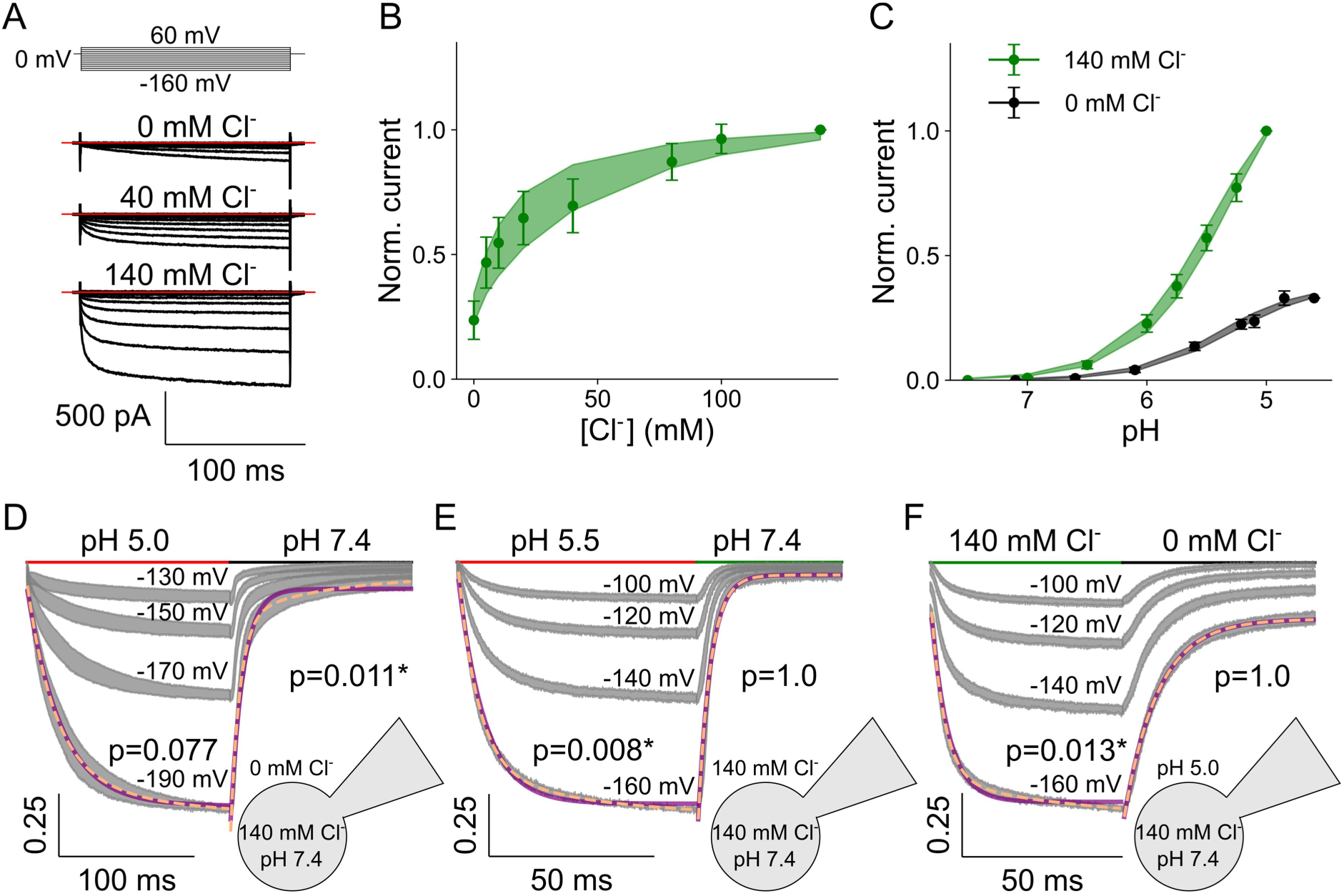
WT VGLUT1_PM_ chloride currents are modulated by voltage, external pH, and [Cl^-^]. Representative WT VGLUT1_PM_ Cl^-^ current responses to voltage steps between -160 mV and +60 mV at pH 5.0 and external [Cl^-^] of 0, 40, and 140 mM (**A**), dose-response plots for VGLUT1_PM_ Cl^-^ currents at pH 5.0 and rising [Cl^-^]_o_ (means ± confidence interval, n = 11 cells, fitted with Michaelis–Menten relationships for a *K*_M_ of 28.3 ± 0.7; **B**); dose-response plots for VGLUT1_PM_ Cl^-^ currents with rising pH at either [Cl^-^]_o_ = 0 (black, means ± confidence interval, n = 10 cells, fitted with Hill relationships for a p*K*_M_ of 5.3 ± 0.003 and a Hill coefficient of 1.2) or 140 mM (green, means ± confidence interval, n = 11 cells, fitted with Hill relationships for a p*K*_M_ of 5.4 ± 0.003 and a Hill coefficient of 1.1; lines and shaded areas depict the mean and 95% confidence interval; **C**), and normalized chloride current responses to pH jumps from pH 7.4 to 5.0 at [Cl^-^]_o_ = 0 mM (**D**), from pH 7.4 to 5.5 at [Cl^-^]_o_ = 140 mM (**E**), or [Cl^-^] jumps from 0 mM to 140 mM and back at pH 5.5 (**F**). Currents were recorded at four continuous voltage levels and shown as 95% confidence interval (gray) from at least 11 experiments, with currents at the most negative voltage fitted with a single (purple line) and double (dashed orange line) exponential functions provided with F-test p-values for each solution change to indicate whether they are better described by biexponential fits, with asterisks marking those that are.

Transporters undergo conformational changes upon ligand association or dissociation, and rates for these processes can be quantified via a kinetic description of transporter currents upon fast substrate application (Grewer, Watzke et al. 2000, Otis and Kavanaugh 2000, Bergles, Tzingounis et al. 2002, Zhang, Tao et al. 2007, Schicker, Uzelac et al. 2012, Ewers, Becher et al. 2013, Kovermann, Hessel et al. 2017, Oh and Boudker 2018, Kortzak, Alleva et al. 2019, Bhat, Niello et al. 2021). Figure 1 D and E depict current responses to pH changes from 7.4 to 5.0 or 5.5 and back to pH 7.4 when the external solution is either Cl^-^ free (Figure 1 D) or has a [Cl^-^]_o_ of 140 mM (Figure 1 E). Fast external solution exchange was achieved by the piezo-driven movement of a perfusion pipette made from dual-channel theta glass tubing (Franke, Hatt et al. 1987, Jonas and Sakmann 1992, Lau, Salazar et al. 2013, Suslova, Kortzak et al. 2023). For clarity, only the four current traces at the most negative voltages are shown. External Cl^-^ accelerates the time course of both processes. Whereas VGLUT1_PM_ Cl^-^ currents activate on monoexponential time courses and deactivate on biexponential time courses upon acidification with a Cl^-^- free external solution, the activation is biexponential and deactivation is monoexponential when the [Cl^-^]_o_ = 140 mM. The application of external Cl^-^ results in biexponential increases and its removal in monoexponential current deactivation (Figure 1 F). Time constants were only minimally voltage dependent and were comparable with and without external Cl^-^. Activation and deactivation time constants were within the same order of magnitude and largely voltage independent (Figure 1— figure supplement 1).

### A kinetic scheme to describe VGLUT1 anion channel function

VGLUT1_PM_ Cl^-^ currents activate and deactivate upon pH jumps between 5.0 and 7.4, with comparable time constants (Figure 1—figure supplement 1). This result excludes the possibility of direct channel activation by H^+^ and indicates that protonated and unprotonated transporters can both open and close, but at distinct rates (Hanke and Miller 1983). A four-state model (with one open and one closed state for each protonation state) predicts monoexponential activation and deactivation upon pH steps. Therefore, we added one doubly protonated open and one doubly protonated closed state to account for biexponential activation or deactivation in response to pH steps. To describe faster activation and deactivation in the presence of external Cl^-^, we assumed that channels with bound Cl^-^ activate or deactivate at different rates and added Cl^-^-bound open and closed states for each protonated state, resulting in a 12-state model (Figure 2). Cl^-^ binding is expected to be diffusion limited, and we combined Cl^-^ binding and associated processes into single rate constants.

**Figure 2:**
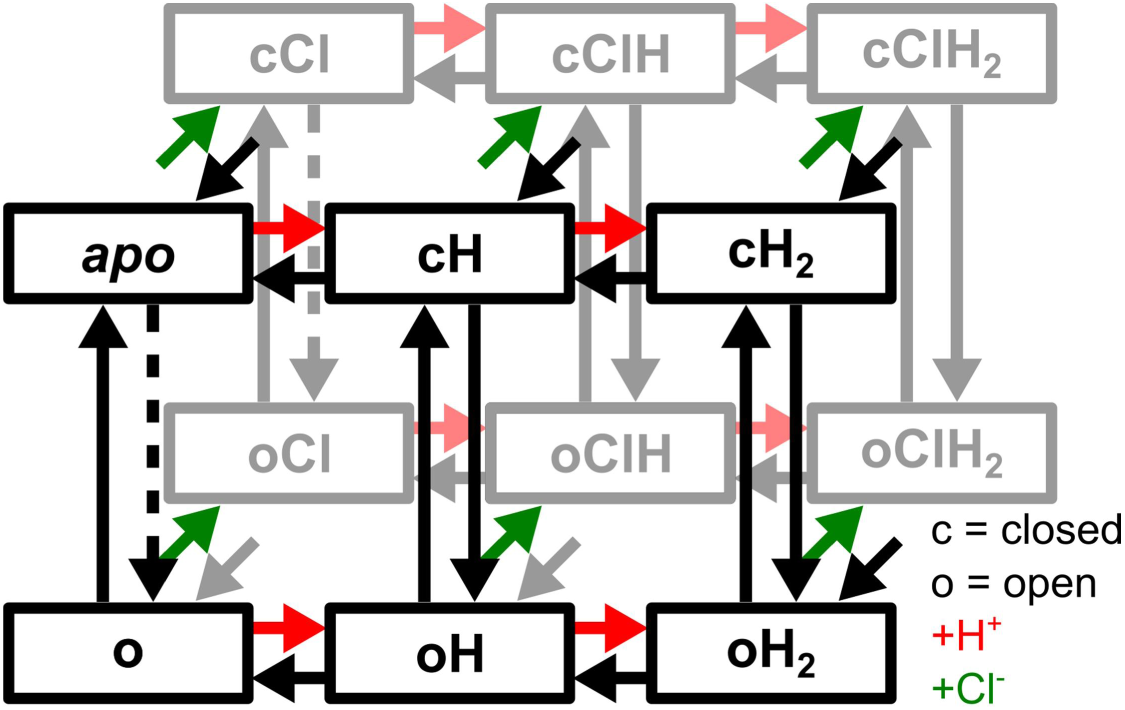
Kinetic model to describe gating of VGLUT1 Cl^-^ channels. Green arrows depict Cl^-^ binding to the black states in front, red arrows depict protonation, and ligand unbinding is shown in corresponding black arrows. The six states on top are *apo* and other closed states, and the channel opens via black downward arrows. Dashed arrows show rate-limited opening without protonation.

This kinetic scheme accurately predicts VGLUT1_PM_ anion channel gating in the presence or absence of external Cl^-^. Figure 3 depicts the fits to normalized current responses from at least 10 HEK293T cells expressing wild-type (WT) VGLUT1_PM_ to voltage steps either from a holding potential of 0 mV (Figure 3 A and D) or after a prepulse to a more negative voltage (Figure 3 B and E), or to pH steps using piezo-driven pipette movements (Figure 3 C and F).

**Figure 3:**
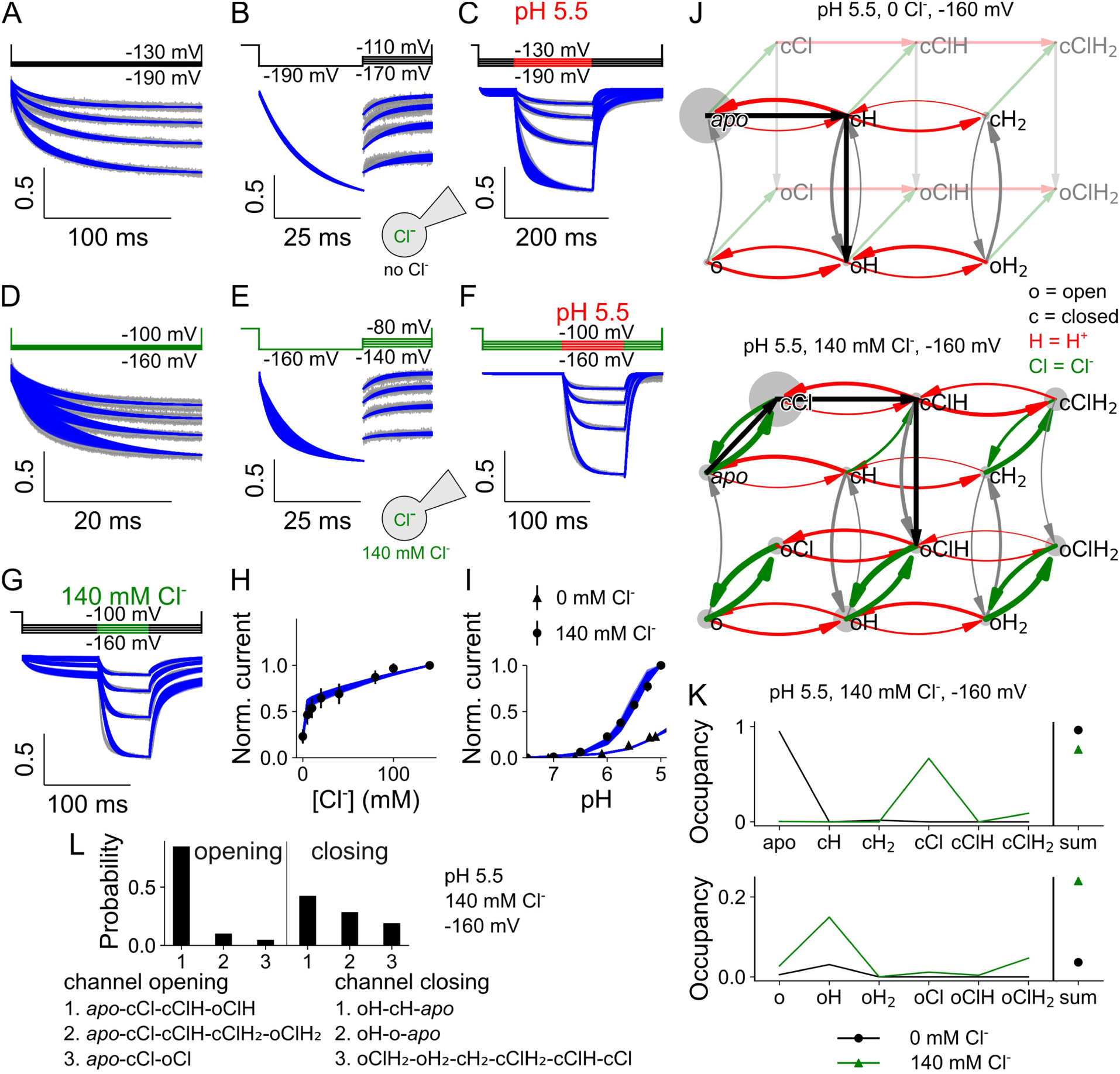
VGLUT1 Cl^-^ currents are well described by a kinetic scheme that assumes three protonation states, each with a closed and open anion channel conformation and all of which can bind Cl^-^. Current responses and predictions from the kinetic scheme for voltage steps from 0 mV to negative potentials with [Cl^-^]_o_ = 0 (A), voltage steps from -190 mV to less negative potentials with [Cl^-^]_o_ = 0 (B), rapid pH steps to pH 5.5 at [Cl^-^]_o_ = 0 (C), voltage steps from 0 mV to negative potentials with [Cl^-^]_o_ = 140 mM (D), voltage steps from -160 mV to less negative potentials with [Cl^-^]_o_ = 140 mM (E), rapid pH steps to pH 5.5 at [Cl^-^]_o_ = 140 mM (F), rapid Cl^-^ steps to 140 mM at pH 5.0 (G), late current plots versus [Cl^-^]_o_ (means ± 95% confidence interval; H), late current plots versus pH at [Cl^-^]_o_ = 0 and 140 mM (I), kinetic schemes describing VGLUT1 Cl^-^ channel activation, with circle size giving occupancy of individual states and curved arrow thickness the rate amplitude for [Cl^-^]_o_ = 0 (top) and [Cl^-^]_o_ = 140 mM (bottom) at pH 5.5 and -160 mV (J), simulated residence probabilities for the indicated closed (top) and open (bottom) states for VGLUT1 anion channels with (green) or without (black) bound Cl^-^ (K), and the three most frequently occurring pathways for activation from *apo* (left) or deactivation from the most common two open states (oH and oClH_2_) to an unprotonated closed state (*apo* or cCl, right; L). Experimental data are given as the 95% confidence interval of patch clamp data (gray) and the fitting results of 250 simulations generated through exploratory mutations (blue) under the same conditions.

Experiments were performed in the absence of external Cl^-^ (Figure 3 A–C) or at [Cl^-^]_o_ = 140 mM (Figure 3 D–F); Figure 3 G shows the fits to current responses upon changes in [Cl^-^]_o_ at an external pH of 5.5, and Figure 3 H and I depict the [Cl^-^] and the pH dependence of normalized late currents. To ensure global model optimization and to assess how well the individual parameters are defined by our experimental data, we randomly modified the fit parameters using an explorative genetic algorithm and collected parameter set values that impaired the goodness of fit by < 25% for each simulated experiment and metric. From these sets of parameters, model predictions from 250 fits are shown as blue lines, evenly spaced from ≥ 10,000 individuals with ≥ 3000 unique values for each variable to provide reliable statistical distinction.

Kinetic modeling enables the calculation and visual tracking of overall fluxes through the states of the kinetic model and, thus, provides insight into activation and deactivation pathways. Figure 3 J shows the kinetic model with the occupancy of individual states depicted as the circle size and rate amplitude as the thickness of curved arrows in the absence or presence of external Cl^-^, and Figure 3 K residence probabilities for the transporter states under both conditions. In the absence of Cl^-^, VGLUT1_PM_ predominantly opens from the singly protonated closed state (as shown in Figure 3 J) and remains singly protonated in the open state (Figure 3 K). Figure 3 L depicts the three most frequent activation and deactivation pathways at pH 5.5, 140 mM Cl^-^, and -160 mV, according to transition path theory-based reactive flux analysis. Under these conditions, VGLUT1_PM_ anion channels predominantly activate via initial Cl^-^ binding, followed by protonation and subsequent channel opening into a singly protonated conformation (oClH). Low-affinity Cl^-^ binding and rapid unbinding make the Cl^-^-free singly protonated state (oH) the most common open state (Figure 3 K). There is a much lower likelihood of channels opening after additional protonation from the doubly protonated Cl^-^ bound open state (oClH_2_). Kinetic modeling also assigns a small probability to WT VGLUT1 opening without protonation (*apo*-cCl-oCl); however, this open state is unstable, leading to negligible channel opening at neutral pH, in agreement with current at pH 7.4 being indistinguishable from background (Kolen, Borghans et al. 2023). Channel deactivation proceeds primarily via three pathways with comparable probabilities: by closing in the singly protonated state, followed by deprotonation; after deprotonation in the open state; or from a doubly protonated Cl^-^-bound state (Figure 3 L).

Figure 4 A depicts the p*K*_a_s for each of the two protonation sites for both Cl^-^-free and Cl^-^-bound VGLUT1, at 0 mV, with open rectangles giving the minimum and maximum of the parameter variants, and at -160 mV, with parameter variants in a violin plot. Negative voltages reduce the *apo* state p*K*_a_ from a median of 2.3 (for a total amplitude range of 2.1–2.4) to 1.1 (0.9–1.2) and after Cl^-^ binding from 4.3 (4.1– 4.5) to 2.0 (1.7–2.1). For the second proton in the closed state, the voltage increases the p*K*_a_ by roughly 2 points, regardless of Cl^-^. This rise, from 6.3 (6.0–6.5) to 8.3 (8.2–8.5) without and from 6.4 (6.1–6.7) to 8.4 (8.1–8.6) with Cl^-^ bound, combined with opening from a single protonated state, explains the relatively high occupancy of cClH_2_ while VGLUT1 is closed (Fig. 3K).

**Figure 4:**
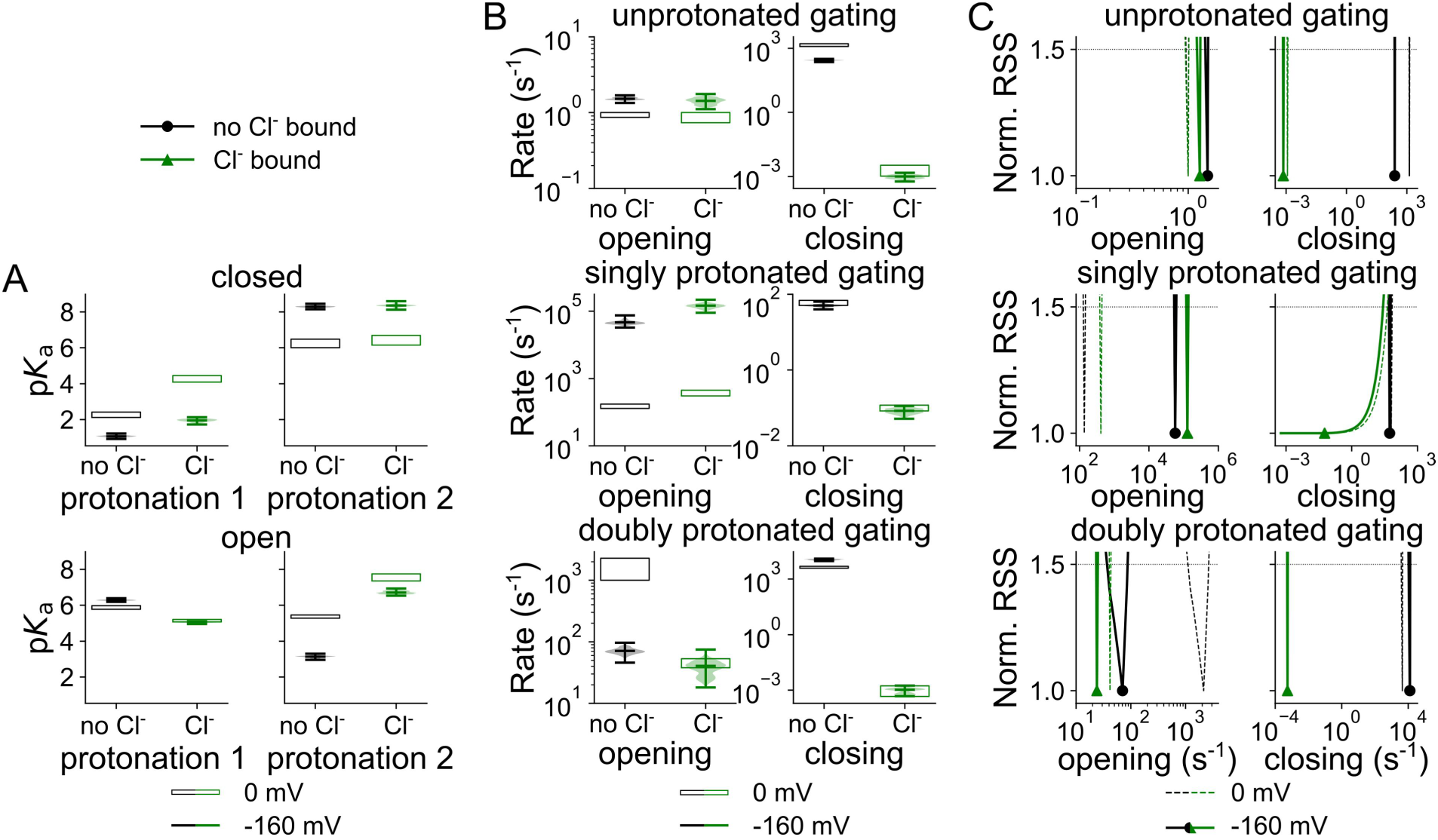
Cl^-^ binding changes VGLUT1 protonation and opening/closing rates. Cl^-^-dependent changes in predicted p*K*_a_s of the first and second protonation sites of the closed (top) or open (bottom) channel at -160 mV (green, Cl^-^ bound; black, no Cl^-^) or at 0 mV (open rectangles or dashed lines; **A**), and violin plots of the amplitude range generated by exploratory mutations (**B**), and rates for channel opening and closing, depicted as normalized RSS representing goodness of fit for a range of amplitudes (**C**).

Figure 4 B and C show opening and closing rates with and without external Cl^-^ at 0 mV or -160 mV. In addition to comparing the full range of parameters obtained in the exploratory fitting (Figure 4 B), we also applied a statistical test to calculate the change in the quality of fit upon only varying the compared parameter (Figure 4 C; Suslova, Kortzak et al. 2023). In such a plot, a sharply defined and distinct parameter value is associated with instantly increasing RSS upon small changes in parameter values without overlap with parameter values obtained under other conditions. The two tests—RSS goodness of fit by parameter amplitude and amplitude distribution via exploratory mutation—demonstrate that—except for closure of the singly protonated channel—all opening and closing transitions are well defined by the fitting procedure and undergo a statistically significant change upon the application of Cl^-^. At -160 mV, Cl^-^ increases opening rates from 4.6×10^4^ (3.3×10^4^–7.5×10^4^, median and parameter amplitude range) s^-1^ to 1.5×10^5^ (9.0×10^4^–2.2×10^5^) s^-1^ in the singly protonated state, but reduces it from 71 (46–97) s^-1^ to 41 (18–75) s^-1^ in the doubly protonated state. The membrane voltage seems to dictate the preference for opening singly protonated, with otherwise low opening rates of 157 (129–174) s^-1^ and 361 (305–451) s^-1^ without and with Cl^-^, respectively. Membrane hyperpolarization decreases the opening rates for doubly protonated VGLUT1 in the absence, but not in the presence of Cl^-^ (Figure 4 B and C).

Cl^-^ reduces the closing rates for all protonation states: at -160 mV from 277 (228– 319) s^-1^ to 9.9×10^-4^ (5.8×10^-4^–1.5×10^-3^) s^-1^ without protonation, from 51 (40–64) s^-1^ to 0.08 (0.05–0.11) s^-1^ for the singly protonated state, and from 1.1×10^4^ (9.4×10^3^– 1.3×10^4^) s^-1^ to 1.1×10^-3^ (4.7×10^-4^–1.8×10^-3^) s^-1^ for the doubly protonated state. The membrane voltage has only minor effects on channel closure.

Cl^-^ binding and unbinding depend on the protonation state and differ for the open and closed conformations (Figure 5, corresponding *z* and *d* parameters are given in Figure 5—figure supplement 1). Cl^-^ association with open channels is close to the limits of diffusion control (10^9^ M^-1^ s^-1^; Wang, Redding et al. 2013). Whereas closed unprotonated and doubly protonated channels bind Cl^-^ at very high rates, at -160 mV with concentration-normalized rates of 6.7×10^8^ (5.8×10^8^–7.2×10^8^) M^-1^ s^-1^ and 1.1×10^7^ (9.4×10^6^–1.3×10^7^) M^-1^ s^-1^, respectively, we observed much lower association rates for the singly protonated closed channel of 1.8×10^3^ (1.6×10^3^–2.1×10^3^) M^-1^ s^-1^. Such association rates may indicate that Cl^-^ binding requires conformational changes that make the binding site accessible in this protonation state. Since changes in the Cl^-^-binding rate will also affect the likelihood of binding-site occupation, it was necessary to distinguish whether protonation adjusts transporter function via altering the binding affinity or association rate for Cl^-^. Therefore, we also assessed how modification of both the association and dissociation rates of Cl^-^ (when held at a fixed ratio to maintain the Cl^-^ dissociation constant, *K*_D_) affect the goodness of fit (Figure 5 B). The analysis demonstrated that only the rate of Cl^-^ association with the singly protonated closed channels determines the fit quality, while for other states such differences could not be demonstrated (Figure 5). Moreover, with constant *K*_D_, only singly protonated Cl^-^ in the closed state is sharply defined, showing the individual association rates are more important here than for any of the other states. We conclude that during anion channel activation VGLUT1 transiently assumes conformational states that are inaccessible to anions in the external solution. Such conformations might be inward facing or occluded with the Cl^-^-binding site inaccessible from either membrane side.

**Figure 5:**
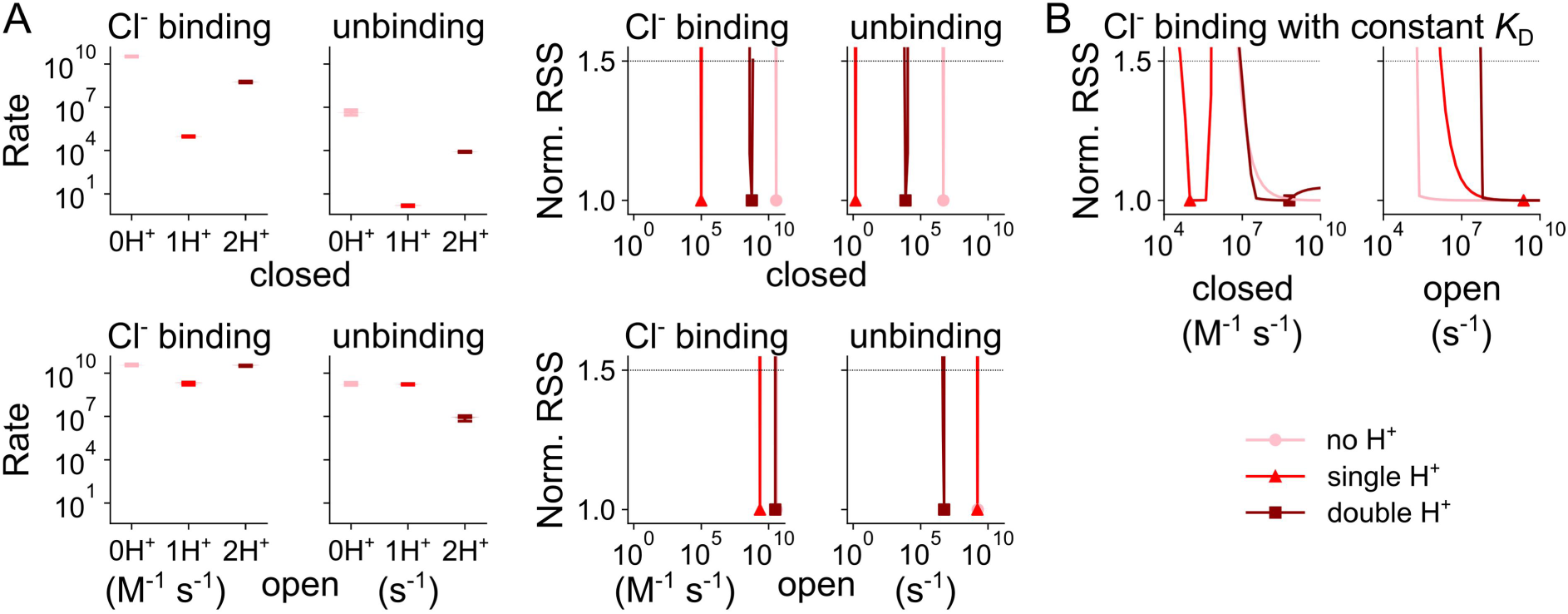
Changes in Cl^-^ association with WT VGLUT1_PM_ with protonation state. Secondary Cl^-^-binding rates to closed (top) or open (bottom) WT VGLUT1_PM_ anion channels in the unprotonated state or the singly or doubly protonated state (light→dark red indicates increasing protonation). Rates are depicted as violin plots (left) showing the amplitude range generated by exploratory mutation and normalized RSS (right) representing goodness of fit for a range of amplitudes (**A**), and Cl^-^ binding at constant *K*_D_, where the unbinding constants were simultaneously altered to maintain the Cl^-^-binding affinity (**B**). All rates are shown at -160 mV.

### H120A impairs Cl^-^ binding to VGLUT1

Substitution of histidine at position 120 by alanine changes the permeation and gating of VGLUT1_PM_ anion channels (Figure 6 A; Kolen, Borghans et al. 2023). We found that the H120A mutation modifies the Cl^-^ dependence of late current amplitudes (H120A: *K*_M_ = 19.4 ± 0.2 mM, WT: *K*_M_ = 28.3 ± 0.7 mM, mean and 95% confidence interval, p = 0.047; Figure 6 B) and changes the p*K*_M_ of acid activation from 5.3 ± 0.003 (WT, Hill coefficient = 1.2) to 5.0 ± 0.004 (H120A, Hill coefficient = 1.2, p = 2.2×10^-6^) in the absence of Cl^-^, but not in the presence of Cl^-^ (5.4 ± 0.003 and Hill coefficient = 1.3, compared to 5.4 ± 0.003 with a Hill coefficient = 1.1 for the WT; Figure 6 C).

**Figure 6:**
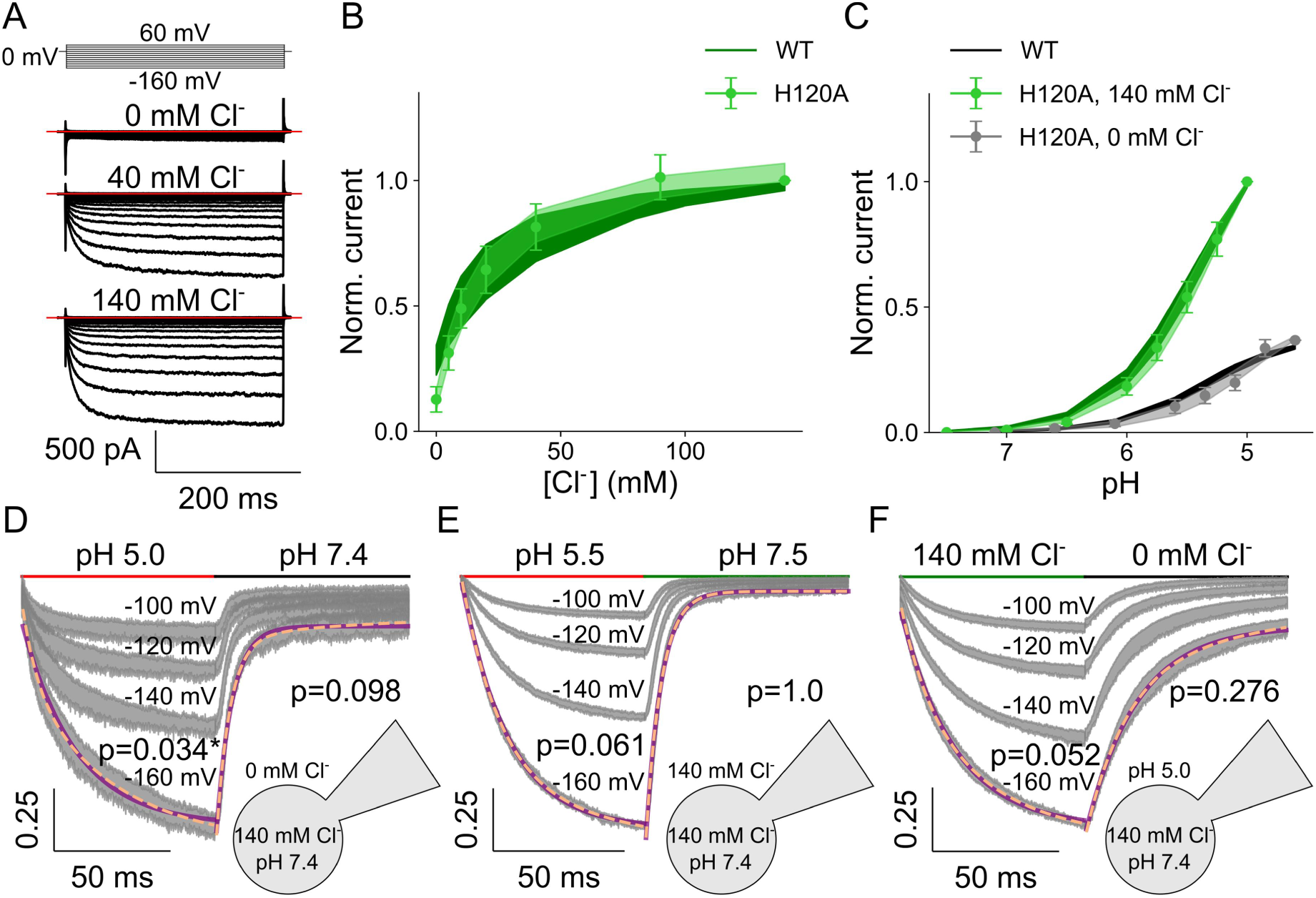
H120A VGLUT1_PM_ chloride currents are modulated by voltage, external pH, and [Cl^-^]. Representative H120A VGLUT1_PM_ Cl^-^ current responses to voltage steps between -160 mV and +60 mV at pH 5.0 and external [Cl^-^] of 0, 40, or 140 mM (**A**), dose-response plots for VGLUT1_PM_ Cl^-^ currents at rising [Cl^-^]_o_ at pH 5.0 (means ± confidence interval, n = 11 cells, fitted with Michaelis–Menten relationships for a *K*_M_ of 19.4 ± 0.2; **B**) or rising pH at [Cl^-^]_o_ = 0 (black, means ± confidence interval, n = 11 cells, fitted with Hill relationships for a p*K*_M_ of 5.0 ± 0.004 and a Hill coefficient of 1.2 for H120A) or 140 mM (green, means ± confidence interval, n = 11 cells, lines and shaded areas depict the mean and 95% confidence interval, fitted with Hill relationships for a p*K*_M_ of 5.4 ± 0.003 and a Hill coefficient of 1.3 for H120A; **C**), and normalized chloride current responses to pH jumps from pH 7.4 to 5.0 at [Cl^-^]_o_ = 0 mM (**D**), or from pH 7.4 to 5.5 at [Cl^-^]_o_ = 140 mM (**E**), or [Cl^-^] jumps from 0 mM to 140 mM and back at pH 5.5 (**F**) held at four continuous voltage levels. Solution change time courses are shown as the 95% confidence interval (gray) from at least 11 experiments; those at the most negative voltage are fitted with a single (purple line) and double (dashed orange line) exponential function and are provided with F-test p-values to indicate whether they are better described by biexponential fits, with asterisks marking those that are.

H120A decelerates VGLUT1_PM_ kinetics in voltage steps (Figure 6 A), pH steps (Figure 6 D and E), and Cl^-^ steps (Figure 6 F). Unlike WT, H120A VGLUT1_PM_ Cl^-^ currents activate and deactivate over a monoexponential time course upon acidification with external Cl^-^. In the absence of external Cl^-^, the kinetics of WT and H120A VGLUT1_PM_ current activation/deactivation upon pH steps are comparable; at high external [Cl^-^], H120A decelerates activation and deactivation (Figure 6—figure supplement 1). Changes in the external [Cl^-^] resulted in monoexponential increases and decreases in the H120A VGLUT1 current (Figure 6 F) at time courses much slower than those of the WT (Figure 6—figure supplement 1); for all conditions, time constants were only minimally voltage dependent.

We reevaluated the parameter amplitudes with the 12 state model developed for WT VGLUT1_PM_, until H120A VGLUT1_PM_ Cl^-^ currents were described well (Figure 7 A–I). Figure 7 J depicts the overall flux through the states of the kinetic model, and Figure 7 K the occupancy of individual states. In the absence of Cl^-^, H120A VGLUT1_PM_ anion channels predominantly open in the singly protonated state, with higher probabilities than the WT; in the presence of Cl^-^, occupation of oClH_2_ was slightly higher for H120A than for WT (Figure 7 K). The H120A mutation alters the activation and deactivation pathways of VGLUT1 channels (Figure 7 L, paths numbered by WT preference). Mutant anion channels primarily open from the doubly protonated state (cClH_2_, path 2), with a lower probability of opening from the singly protonated Cl^-^-bound state. Whereas the WT mainly closes from the oH state, either by direct deprotonation or by deprotonation of the closed state, H120A VGLUT1_PM_ mainly passes through Cl^-^ unbinding to oH_2_ and then to cH_2_, cClH_2_, cClH, and, finally, to cCl (path 3).

**Figure 7:**
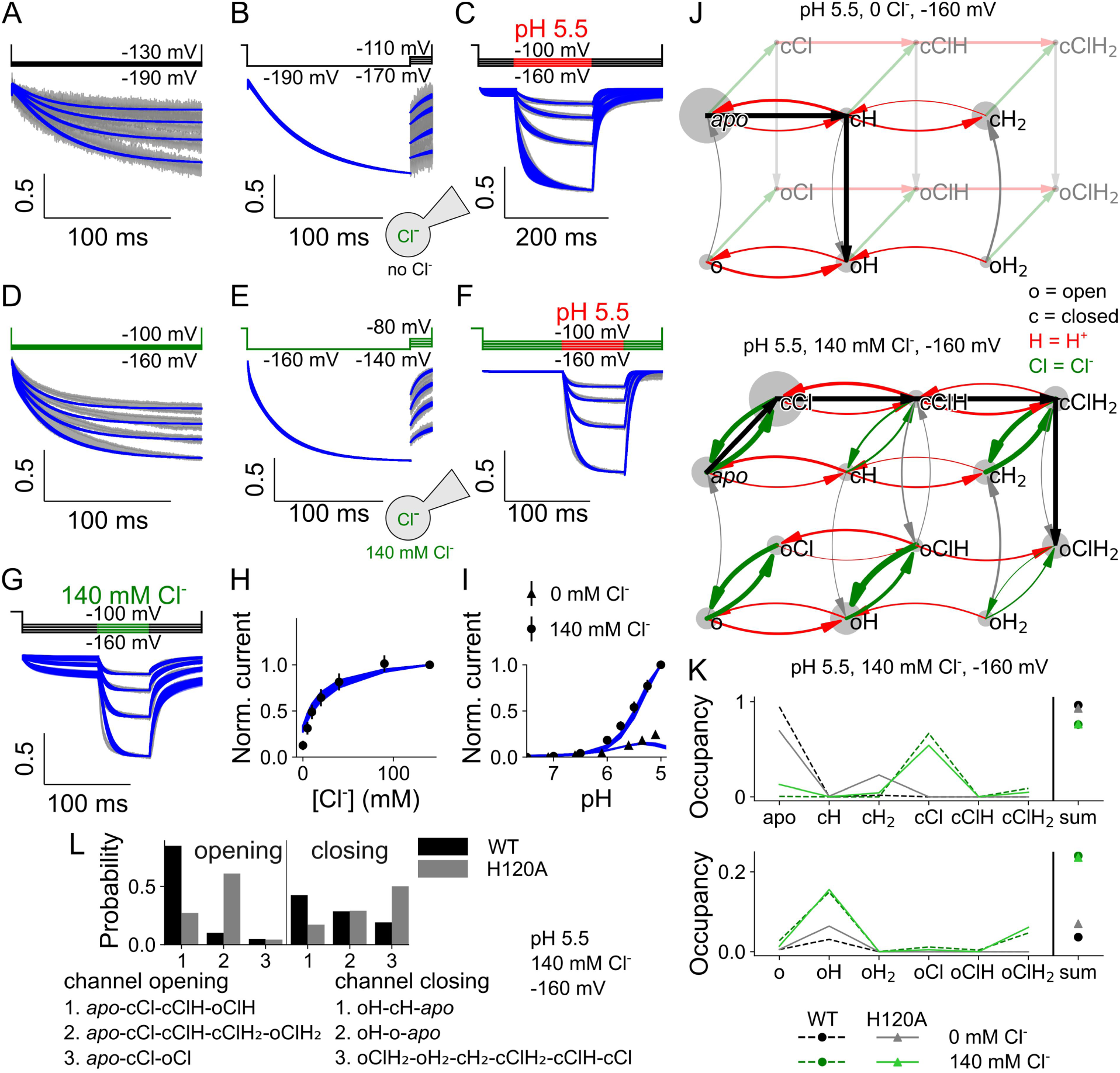
H120A VGLUT1 Cl^-^ currents are well described by a kinetic scheme that assumes three protonation states, each with a closed and open anion channel conformation and all of which can bind Cl^-^. Current responses and predictions by the kinetic scheme for voltage steps from 0 mV to negative potentials at [Cl^-^]_o_ = 0 mM (A), more negative to less negative potentials at [Cl^-^]_o_ = 0 (B), pH steps from pH 7.4 to pH 5.5 at [Cl^-^]_o_ = 0 (C), voltage steps from 0 mV to negative potentials at [Cl^-^]_o_ = 140 mM (D), more negative to less negative potentials at [Cl^-^]_o_ = 140 mM (E), pH steps from pH 7.4 to pH 5.5 at [Cl^-^]_o_ = 140 mM (F) or upon changes in [Cl^-^]_o_ from 0 to 140 mM at pH 5 (G), plots of late currents (means ± 95% confidence interval) versus [Cl^-^]_o_ (H) or pH (I), kinetic schemes describing H120A VGLUT1 Cl^-^ channel activation for [Cl^-^]_o_ = 0 mM (top) and [Cl^-^]_o_ = 140 mM (bottom, with circle size giving occupancy of individual states and curved arrow thickness the rate amplitude; J), simulated residence probabilities for the indicated closed (top) and open (bottom) states for H120A VGLUT1 anion channels with (green) or without (black) bound Cl^-^ (K), and the three most frequently occurring activation (left) or deactivation (right) pathways upon a rapid pH change (L). Experimental data are given as the 95% confidence interval of patch clamp data (gray) and the fitting results of 250 simulations from parameter sets generated through exploratory mutations (blue) under the same conditions.

Without Cl^-^, H120A strongly increases the p*K*_a_ of the first protonation site of the closed channel from 1.1 (0.9–1.2), median and parameter amplitude range) to 3.7 (3.5–3.9; all values obtained at -160 mV; Figure 8 A). Cl^-^ binding reduces this value back to 3.1 (2.8–3.2). H120A impairs protonation of the second site, without effect of Cl^-^. For Cl^-^-bound open VGLUT1, H120A decreases the p*K*_a_ of the first protonation site from 5.0 (5.0–5.1) to 2.7 (2.6–3.0), with a smaller increase from 6.7 (6.5–6.9) to 7.8 (7.6–8.0) for the second protonation site. H120A decreases opening rates for singly protonated states, from 4.6×10^4^ (3.3×10^4^–7.5×10^4^) s^-1^ to 74 (55–93) s^-1^ without Cl^-^ and from 1.5×10^5^ (9.0×10^4^–2.2×10^5^) s^-1^ to 1439 (1044–1807) s^-1^ with Cl^-^ (Figure 8 B). It increases closing rates in the presence of Cl^-^: from 9.9×10^-4^ (5.8×10^-4^–1.5×10^-3^) s^-1^ to 0.026 (0.012–0.050) s^-1^ when unprotonated, from 1.1×10^-3^ (4.7×10^-4^–1.8×10^-3^) s^-1^ to 1.0 (0.5–1.5) s^-1^ when doubly protonated; with single protonation, the rates are not sharply defined. Moreover, for doubly protonated states, H120A has a lower opening rate than the WT in the absence of Cl^-^ with 4.2 (2.7–5.7) s^-1^ versus 71 (46– 97) s^-1^, but a higher rate in its presence with 250 (85–532) s^-1^ versus 41 (18–75) s^-1^. This likely forms the basis for its preferential channel opening under double protonation only in the presence of Cl^-^.

**Figure 8:**
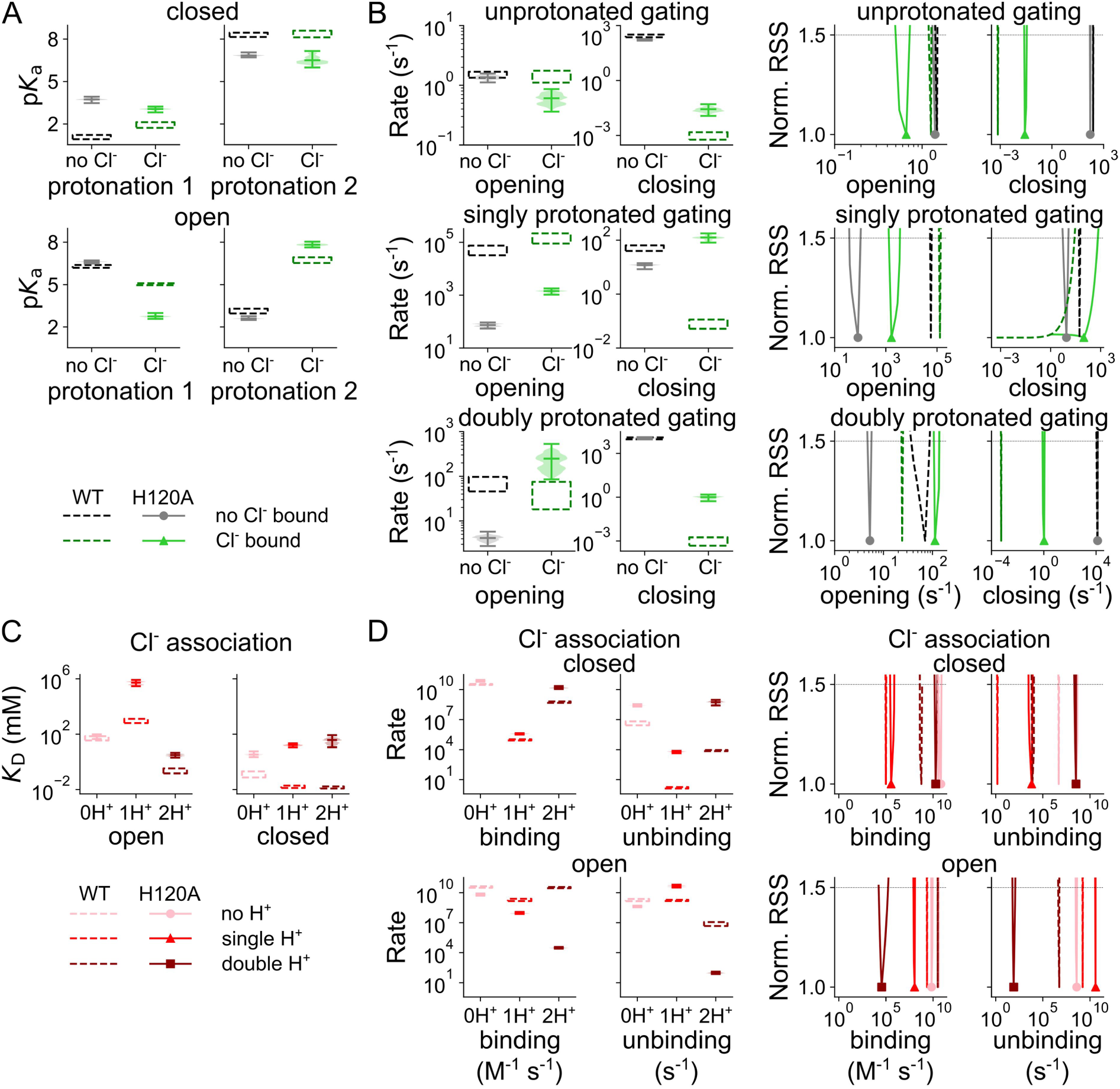
Cl^-^ binding changes H120A VGLUT1_PM_ protonation and opening/closing rates, protonation changes Cl^-^ binding. Cl^-^-dependent changes in p*K*_a_ for the first and second protonation sites of the closed (top) or open (bottom) H120A VGLUT1_PM_ channels (**A**), opening and closing rates (no Cl^-^ in black, Cl^-^-bound in green, WT in darker dashed lines) for different protonation states (**B**), Cl^-^ association *K*_D_ at different protonation states (light→dark red indicates increasing protonation; **C**), and Cl^-^ binding and unbinding rates for closed and open H120A VGLUT1_PM_ channels at different protonation states (**D**). Rates are depicted as violin plots (left) showing the amplitude range generated by exploratory mutation and normalized RSS (right) representing goodness of fit for a range of amplitudes. All rates are shown at -160 mV.

H120A alters the preferred pathways of channel activation through changes in channel opening and closing, effectively switching its primary Cl^-^-activated opening state from single to double protonation. Figure 8—figure supplement 1 depicts the effects of the H120A mutation on the voltage dependence of protonation, opening/closing, and Cl^-^ association in the absence and presence of luminal/external Cl^-^.

The H120A mutation has pronounced effects on Cl^-^ binding and unbinding (Figure 8 C). Whereas H120A left Cl^-^ dissociation constants for the unprotonated open channel unaffected, it increased the *K*_D_ under all other conditions, in the open channel from 869 (641–1263) mM to 5.8×10^5^ (2.9×10^5^–8.3×10^5^) mM when singly protonated, and from 0.24 (0.15–0.36) mM to 3.1 (2.0–4.5) mM when doubly protonated. For the closed channel, dissociation constants increase from 0.12 (0.08– 0.20) mM to 3.4 (2.2–5.8) mM when unprotonated, from 0.017 (0.013–0.019) mM to 16 (11–22) mM when singly protonated, and from 0.014 (0.012–0.017 mM to 38 (11– 85) mM when doubly protonated. These changes correspond to altered Cl^-^ association rates (Figure 8 C). However, by testing the variation in Cl^-^ binding to open or closed channel at a fixed affinity (Figure 8—figure supplement 2), we demonstrate that H120A does not affect the accessibility but, rather, the affinity of the binding sites.

### A kinetic scheme to describe VGLUT1 glutamate and aspartate transport

VGLUT1_PM_ glutamate or aspartate currents are much smaller than Cl^-^ currents, requiring complete substitution of intracellular Cl^-^ with these anions to record such transport currents (Kolen, Borghans et al. 2023). Figure 9 depicts representative glutamate and aspartate currents, as well as the pH and [Cl^-^] dependence of late currents at -160 mV, with Figure 9—figure supplement 1 showing the time constants of pH and [Cl^-^] jumps. Aspartate currents differ from glutamate, with a slightly higher amplitude and a more negative current reversal potential (Figure 9 A and B), reflecting the difference in transport stoichiometry. Glutamate and aspartate currents are close to background in the absence of Cl^-^; therefore, we only recorded current responses to pH steps at [Cl^-^]_o_ = 40 mM (Figure 9 C and D) or to [Cl^-^] steps at acidic pH (Figure 9 E and F). For glutamate and aspartate, pH jumps resulted in monoexponential activation or deactivation, with closely similar kinetics (Figure 9 C and D). The pH dependence of late glutamate and aspartate currents can be fitted with the Hill equation, and the p*K*_M_ values for both are indistinguishable (glutamate: 5.5 ± 0.005 with a Hill coefficient of 1.3, aspartate: 5.5 ± 0.002 with a Hill coefficient of 1.4, p = 0.53, mean and 95% confidence interval; Figure 9 G) and similar to the values obtained with Cl^-^ currents.

**Figure 9:**
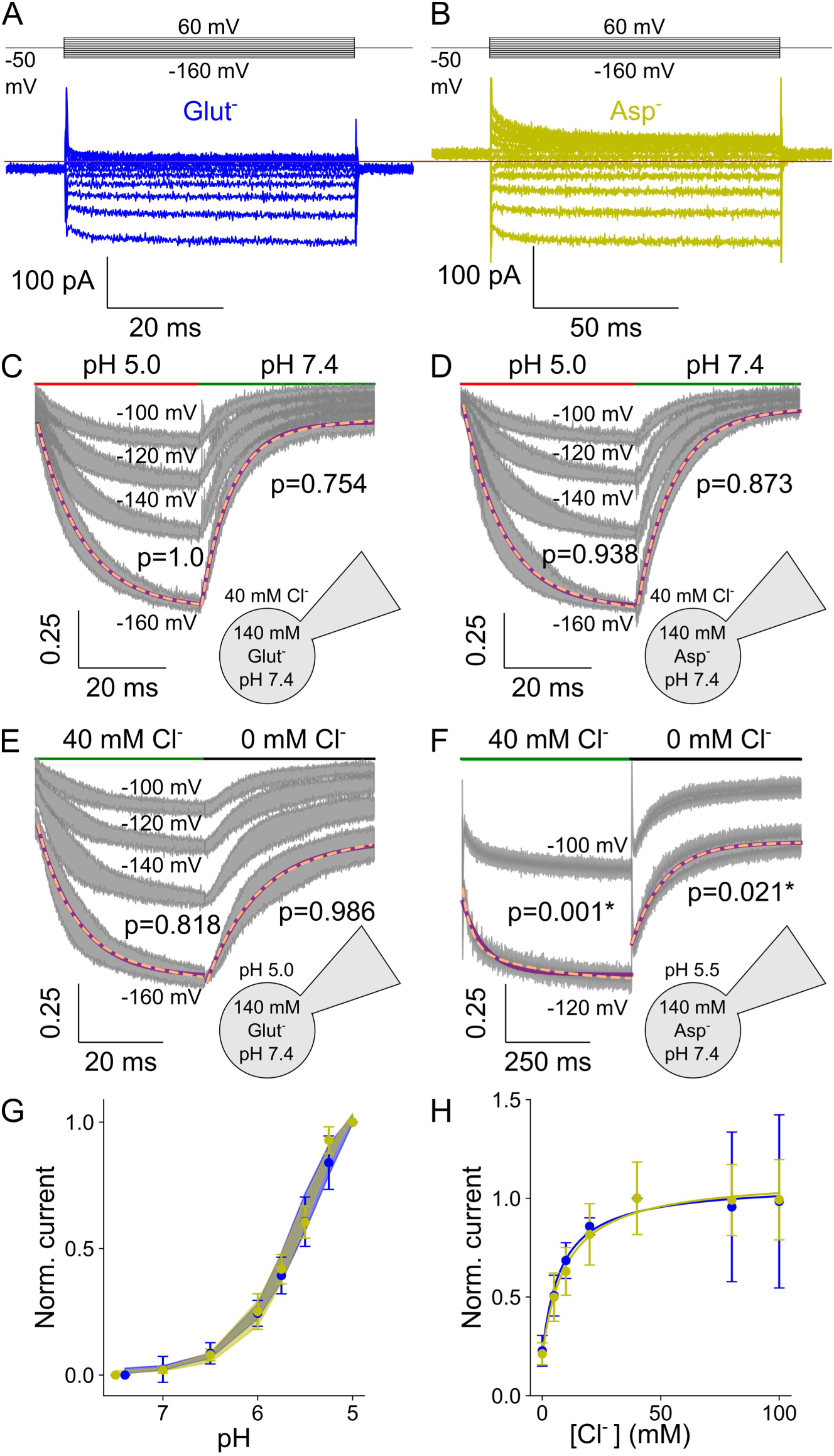
VGLUT1PM glutamate and aspartate current are modulated by voltage, external pH, and [Cl^-^]. Representative current responses of transfected cells dialyzed with glutamate-based (blue; **A**) or aspartate-based (yellow; **B**) solutions to voltage steps with [Cl^-^]_o_ of 40 mM and the pH of 5.5, current responses to pH steps from pH 7.4 to pH 5.0 with glutamate (**C**) or aspartate (**D)**, or to concentration steps from [Cl^-^]_o_ = 0 mM to 40 mM with glutamate (**E**) or aspartate (**F**). pH dependence of normalized steady-state current amplitudes at [Cl^-^]_o_ = 140 mM and a membrane potential of -160 mV bootstrapped using a Hill relationship with glutamate (p*K*_M_ = 5.5 ± 0.005, Hill coefficient = 1.3) and aspartate (p*K*_M_ = 5.5 ± 0.002, Hill coefficient = 1.4, [Cl^-^]_o_ = 40 mM; **G**), and the [Cl^-^] dependence of steady-state current means at pH 5.0 and -160 mV fitted to a Michaelis–Menten relationship with glutamate (*K*_M_ = 8.1 ± 2.0 mM) and aspartate (*K*_M_ = 9.9 ± 2.2 mM; pH 5.5, -140 mV; **H**). Currents (gray) and error bars represent the 95% confidence interval from ≥ 10 measurements; currents at the most negative voltage are fitted with a single (purple line) and double (dashed orange line) exponential function along with F-test p-values to indicate whether they are better described by biexponential fits, with asterisks marking those that are.

Cl^-^ concentration jumps elicited very different responses for the glutamate and aspartate currents. Rapid increases in [Cl^-^]_o_ from 0 to 40 mM resulted in monoexponential increases of glutamate currents with time constants of around 10 ms, and the currents decayed upon decreasing steps to [Cl^-^] of 0 mM over a similar time course (Figure 9). For aspartate currents, Cl^-^-dependent activation was more than 10-fold slower, and followed a biexponential time course upon [Cl^-^] increases or decreases with time constants of around 140 ms (Figure 9 F, Figure 9—figure supplement 1 B). Despite these differences in kinetics, glutamate and aspartate currents rose with increasing external [Cl^-^] with a similar concentration dependence (i.e. glutamate: K*_M_* = 8.1 ± 2.0 mM, aspartate: 9.9 ± 2.2 mM, mean and 95% confidence interval; Figure 9 H). These values are significantly lower than the *K*_M_ of 28.3 ± 0.7 mM for VGLUT1 Cl^-^ currents (p = 0.0097, n ≥ 10 cells, one-sample Wilcoxon signed rank test between the combined glutamate mean fit and individual cellular fits of Cl^-^ current).

We calculated unitary aspartate transport rates from combined whole-cell current and fluorescence amplitudes of an earlier publication: in this dataset, whole-cell current amplitudes increase linearly with cellular expression levels quantified as whole-cell fluorescence, and we compared fluorescence-normalized current values from this publication in combination with its calculated glutamate transport of 561 ± 123 s^-1^ (Kolen, Borghans et al. 2023). Since aspartate currents exceed glutamate by a factor 2.3 at -160 mV, without the protons exchanged with glutamate at a stoichiometry of 1:1, we obtained a transport rate of approximately 2581 s^-1^.

VGLUT1 structurally resembles transporters of the Major Facilitator Superfamily (Li, Eriksen et al. 2020) and operates as a secondary active glutamate transporter. Therefore, we described glutamate and aspartate transport with an alternating access mechanism that shuttles between inward-facing states that permit the binding of amino acids from the cytoplasm and outward-facing states that release amino acids to the vesicular lumen (Figure 10).

**Figure 10:**
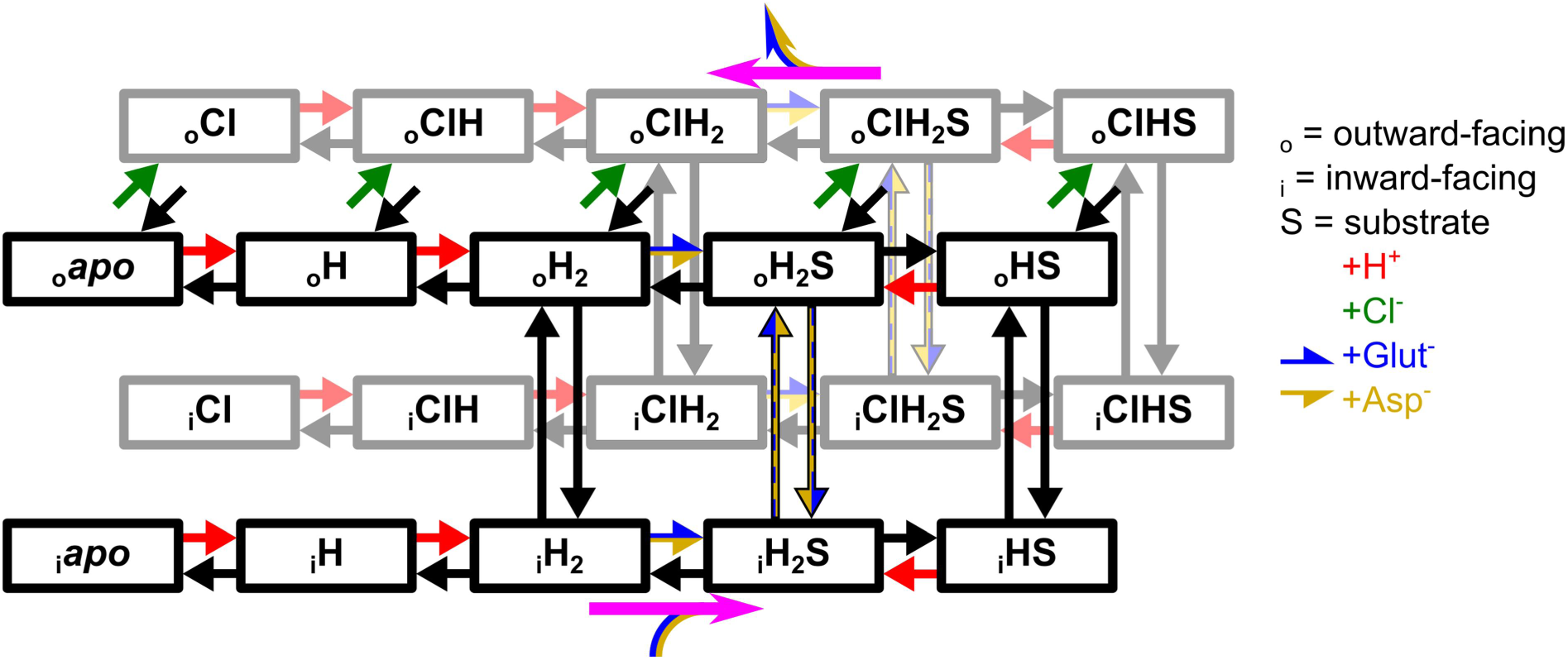
Kinetic model of active glutamate and aspartate transport by VGLUT1. The top states are outward-facing, connected to the inward-facing states at the bottom by vertical arrows. Green arrows depict Cl^-^ binding to the (outward-facing) black states in front, red arrows depict protonation, substrate binding is shown in horizontal blue-yellow arrows, and ligand unbinding in corresponding black arrows. Vertical black arrows with a blue-yellow fill show doubly protonated translocation with substrate, with limited rates when glutamate-bound. Pink arrows indicate the general direction of the Cl^-^-free (black states) and Cl^-^-bound (gray states) transport cycle.

The model is based on multiple assumptions. Since the Cl^-^ channel is described using two protonation sites and shows a pH dependence remarkably similar to VGLUT1 transport of glutamate and aspartate, we assumed that the same number of sites are protonated when the empty transporter returns to the inward-facing conformation. Substrates then bind to the doubly protonated transporter and cross the membrane either in this protonation state or after the release of one proton. Substrate translocation in the doubly protonated state results in uncoupled uniport, and singly protonated transitions in H^+^-substrate exchange. To permit H^+^ transport, transporter protonation and deprotonation were made possible in the inward- and outward-facing conformations. To ensure stoichiometrically coupled H^+^-glutamate exchange, H^+^ fluxes that are not coupled to glutamate transfer need to be prevented because they enable passive H^+^ flux. Therefore, for transporters with no substrate bound, we permitted only doubly protonated translocation.

Cl^-^ is only present in the external solution, and we therefore exclusively accounted for Cl^-^ binding in the outward-facing conformation. Since we studied glutamate and aspartate currents at negative voltages in the absence of internal Cl^-^, anion-conducting states were not added to the kinetic scheme. The scheme correctly describes the current kinetics and pH and [Cl^-^] dependence of both glutamate and aspartate currents (Figure 11), with both H^+^-glutamate and aspartate transport rates similar to experimental data (Kolen, Borghans et al. 2023).

**Figure 11:**
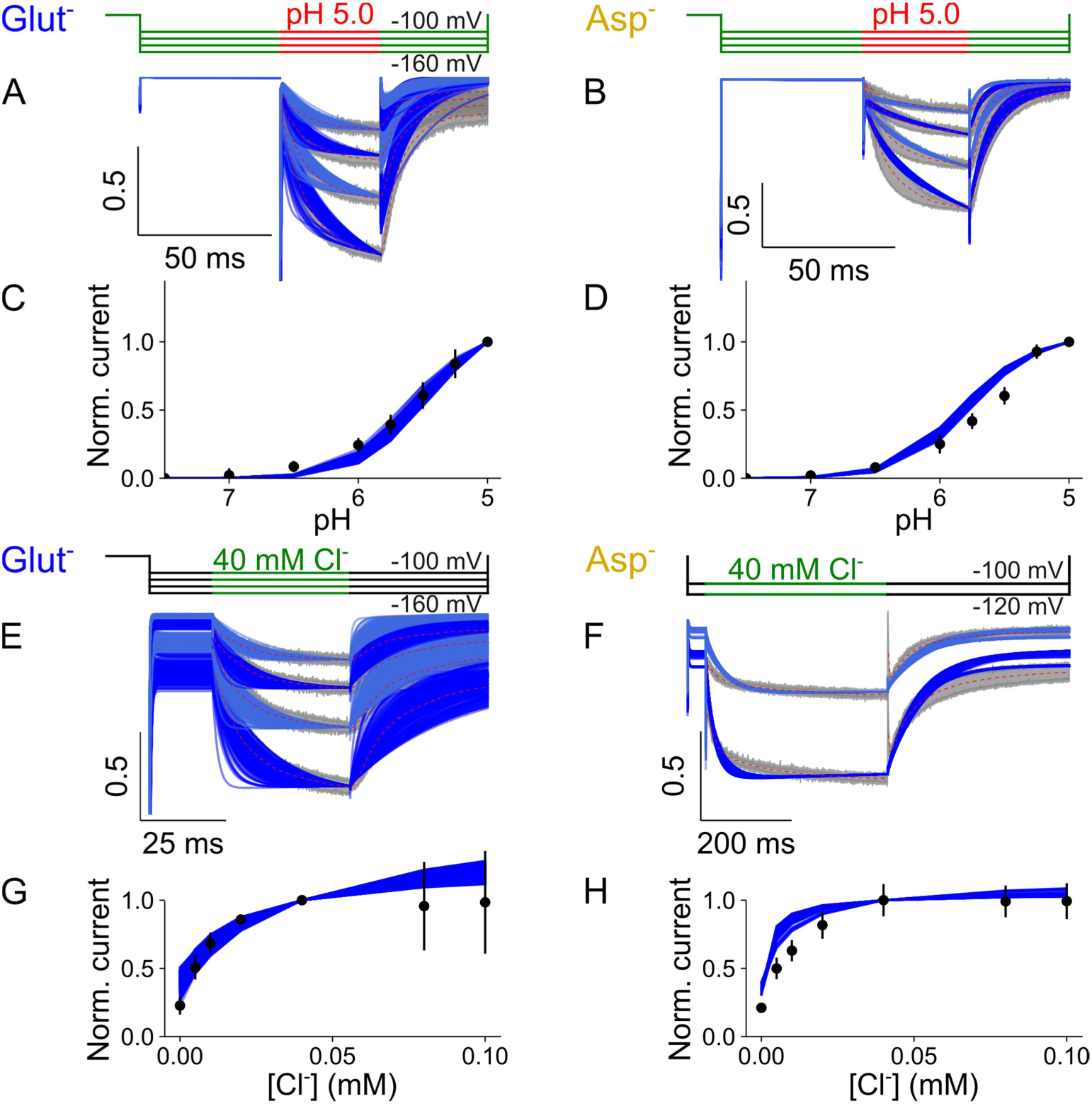
VGLUT1PM glutamate and aspartate currents are well described by an alternating access kinetic scheme. Experimental currents and predictions by the kinetic scheme for pH steps from 7.4 to 5.0 at [Cl^-^]_o_ = 140 mM with glutamate (A) or aspartate (B), plots showing late currents (means ± 95% confidence interval) against pH for glutamate (C) or aspartate (D), experimental currents and predictions by the kinetic scheme for changes in [Cl^-^]_o_ from 0 to 40 mM at pH 5.0 for glutamate (E) or at pH 5.5 for aspartate (F), and plots showing late currents (means ± 95% confidence interval) against pH at [Cl^-^]_o_ = 40 mM for glutamate (G) or aspartate (H). All panels show experimental data as the 95% confidence interval of patch clamp data (gray and black) and the fitting results of 250 simulations from parameter sets generated through exploratory mutations (blue) under the same conditions.

### Cl^-^ stimulates active transport by modifying glutamate and proton binding

Figure 12 compares the kinetic parameters for coupled glutamate transport in the Cl^-^- free and Cl^-^-bound states; its voltage dependence is given by the corresponding *z* and *d* parameters in Figure 12—figure supplement 1. VGLUT1 binds substrates from the internal solution after double protonation and translocation to the inward-facing state. At -160 mV, external Cl^-^ enhances the rates of glutamate binding to the inward-facing state from 6.9×10^4^ (3.5×10^4^–9.1×10^4^, median and parameter amplitude range) M^-1^ s^-1^ to 6.0×10^6^ (2.8×10^6^–2.4×10^7^) M^-1^ s^-1^ (Figure 12 B). Changing the glutamate binding rate while modifying unbinding to maintain the same *K*_D_ results in sharply defined and distinct RSS values in both conformations, which shows that Cl^-^ increases individual association rates and not just the *K*_D_ (Figure 12—figure supplement 2). This change in association rates suggests that an occluded glutamate-binding pocket can open to the cytoplasm upon Cl^-^ binding. Cl^-^ does not affect predicted p*K*_a_ values for protonation after glutamate binding, they are close to pH 1 without Cl^-^ at 1.7 (0.6–2.1) or with Cl^-^ at 0.3 (-0.6–1.3). These low p*K*_a_ values ensure deprotonation of the inward-facing glutamate-bound transporter at neutral cytoplasmic pH (Figure 12 C).

**Figure 12:**
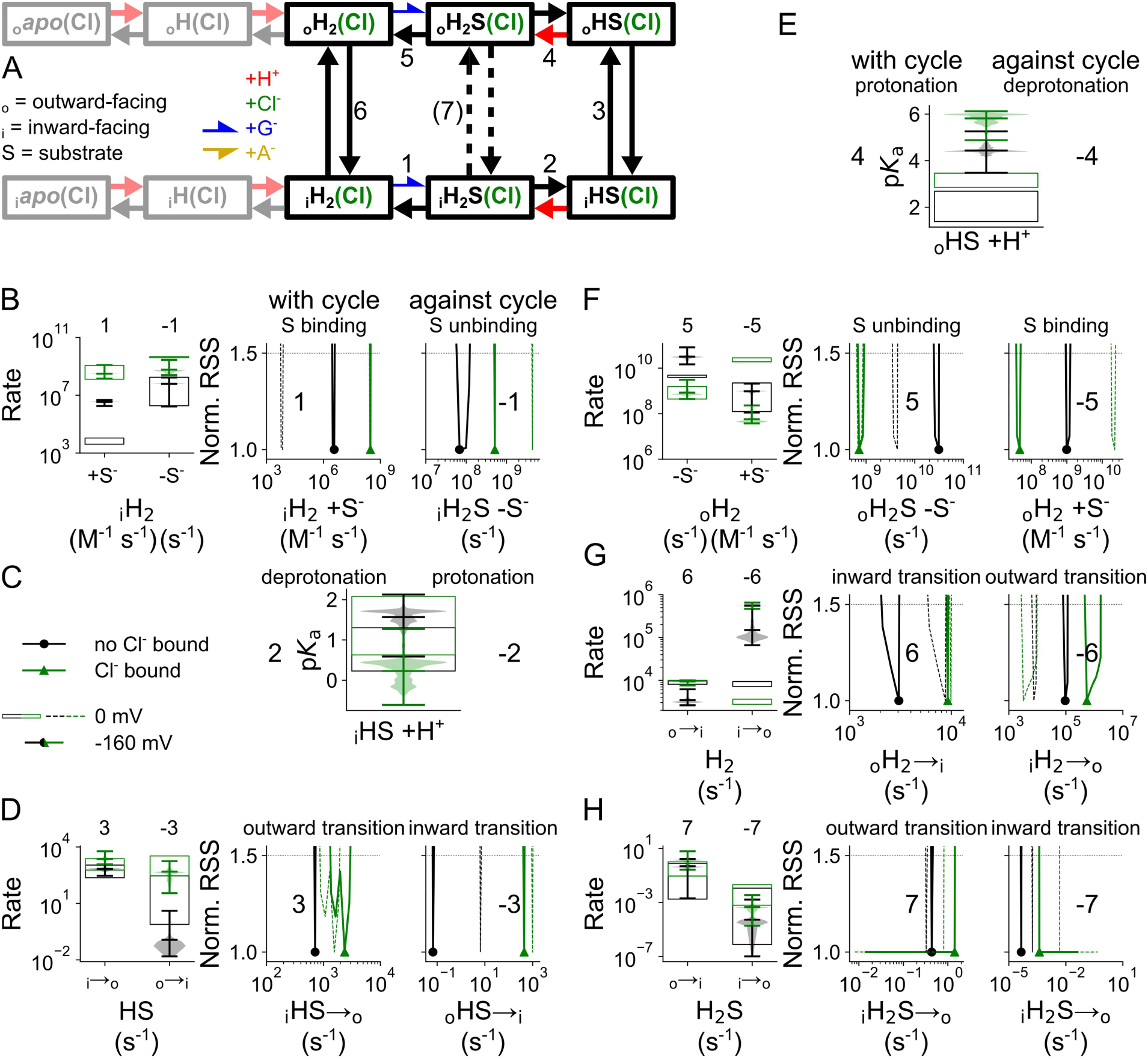
Cl^-^ binding changes VGLUT1 glutamate transport parameters. Kinetic scheme describing glutamate transport by VGLUT1 with or without bound Cl^-^ (A), Cl^-^- induced changes in glutamate binding/unbinding in the inward-facing conformation (B), in the p*K*_a_ of the second protonation site in the inward-facing conformation (C), in the singly protonated translocation rate (D), in p*K*_a_ for the second protonation site in the outward-facing conformation (E), in glutamate binding/unbinding in the outward-facing conformation (F), and in the doubly protonated translocation rate without substrate (G) or doubly protonated translocation rate with substrate (H). Rates along the transport cycle are positively numbered and on the left, next to rates going against the transport cycle with negative numbers. Protonation is represented by p*K*_a_ violin plots; other simulated rates are given as normalized RSS representing goodness of fit for a range of amplitudes in addition to violin plots depicting the amplitude range generated by exploratory mutation. All rates are shown at -160 mV or at 0 V in open rectangles or dashed lines.

Outward translocation rates of transporters with glutamate bound as substrate (_i_HS) did not differ with and without Cl^-^ (Figure 12 D), and neither did the p*K*_a_ of the subsequent protonation (Figure 12 E). Glutamate release to the lumen was impaired by Cl^-^, from 3.1×10^10^ (1.5×10^10^–8.1×10^10^) s^-1^ to 7.4×10^8^ (4.4×10^8^–3.1×10^9^) s^-1^ (Figure 12 F), which fits with Cl^-^ causing faster glutamate binding in the inward-facing conformation. With a small increase in the final substrate-free (_o_H_2_) inward translocation step from 3152 (2608–6165) to 9293 (7602–9966) s^-1^ (Figure 12 G), Cl^-^ provides a net positive effect on glutamate transport. We set an upper limit of 1 s^-1^ to the translocation rate constant of the doubly protonated substrate-bound transporter (_o_H_2_S) to the inward-facing state only with glutamate as the substrate (Figure 12 H).

Glutamate transport is pronouncedly voltage dependent. The comparison of kinetic parameters at 0 mV and -160 mV in the presence of Cl^-^ demonstrate that the main effect of voltage is stimulation of the protonation of the substrate bound transporter.

### Allosteric regulation of transporter protonation contributes to VGLUT1 substrate selectivity

Figure 13 compares the p*K*_a_ values and transition rates for glutamate and aspartate transport in the Cl^-^-bound cycle, its voltage dependence is given by the corresponding *z* and *d* parameters in Figure 13—figure supplement 1. Glutamate and aspartate bind at comparable rates to the doubly protonated inward-facing transporter, but with 5.1×10^8^ (2.5×10^8^–2.8×10^9^, median and parameter amplitude range at -160 mV) s^-1^ glutamate unbinding is significantly faster than aspartate unbinding with 309 (181–366) s^-1^ (Figure 13 A and B). The p*K*_a_ of the second protonation site is 0.3 (-0.6–1.3) for glutamate-bound transporters and 5.1 (5.0–5.4) for aspartate-bound transporters (Figure 13 C). This difference results in deprotonation of the glutamate-bound, but not of the aspartate-bound transporter at neutral cytoplasmic pH, and supports glutamate translocation in the singly protonated state and H^+^-glutamate exchange. We tested the importance of the different substrate-bound p*K*_a_ values by modifying unbinding rates in both inward- and outward-facing conformation. Increasing the low glutamate value and decreasing the high aspartate value reduces transport rates, showing how important the optimized rates are for VGLUT1 transport rates (Figure 13—figure supplement 2).

**Figure 13:**
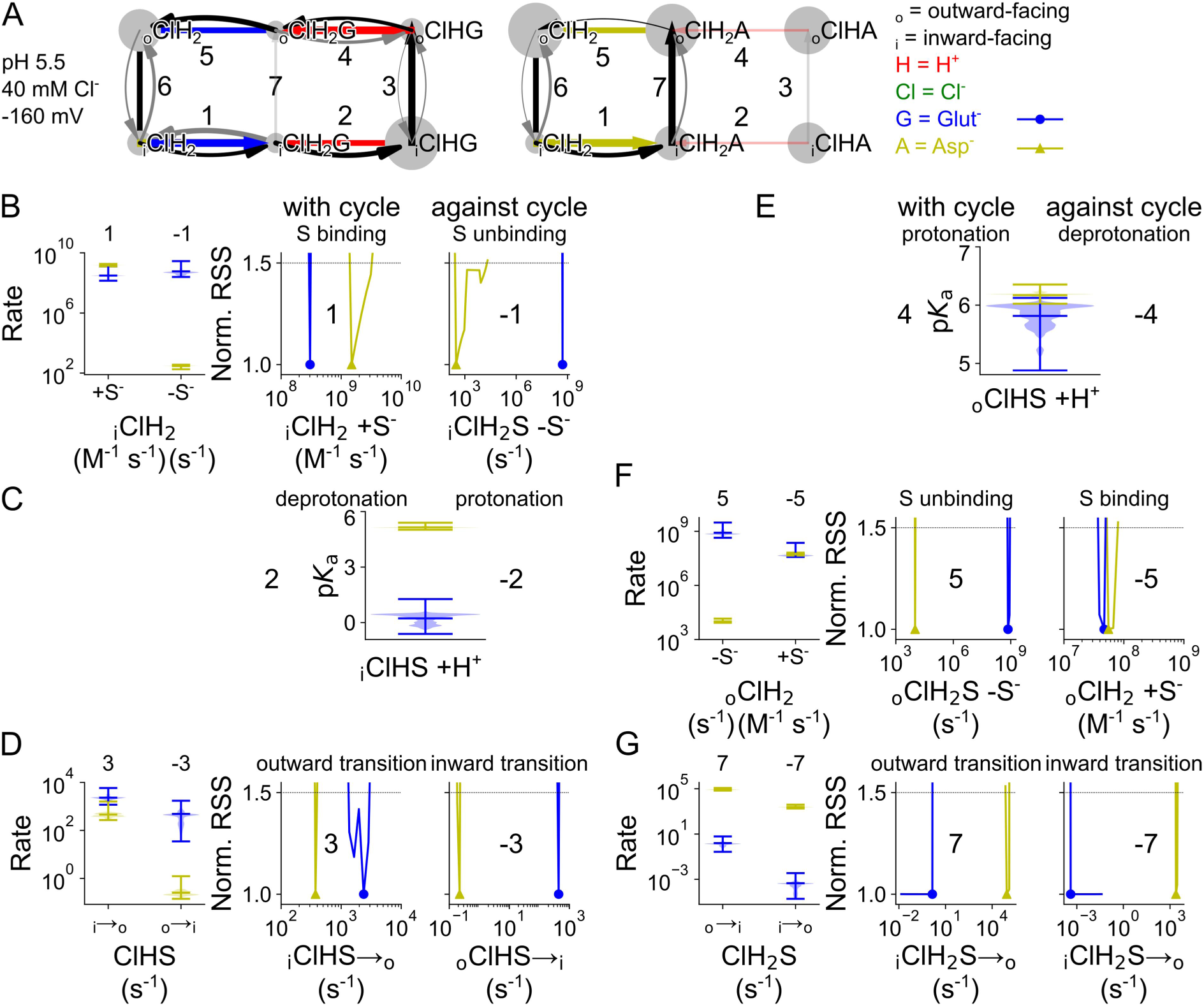
Differences in the VGLUT1 glutamate and aspartate transport cycle. Kinetic scheme describing Cl^-^-bound and protonated VGLUT1 glutamate (left) or aspartate (right) transport function (**A**), rates of substrate binding/unbinding in the inward-facing conformation (**B**), p*K*_a_ values for the second protonation site in the inward-facing conformation (**C**), translocation rates for the singly protonated, substrate-bound conformation (**D**), p*K*_a_ values for the second protonation site in the outward-facing conformation (**E**), rates of substrate binding/unbinding in the outward-facing conformation (**F**), and translocation rates for the doubly protonated, substrate-bound conformation (**G**). Protonation is described by p*K*_a_ violin plots; other simulated rates are given as normalized RSS representing goodness of fit for a range of amplitudes in addition to violin plots depicting the amplitude range generated by exploratory mutation. Data for glutamate transport are shown in blue and for aspartate transport in yellow. All rates are shown at -160 mV.

Substrate-bound outward translocation in the singly protonated state is the same for glutamate and aspartate (Figure 13 D). Outward-facing p*K*_a_ values with aspartate were the same as the glutamate-bound transporter (Figure 13 E), at a value comparable to the inward-facing aspartate-bound p*K*_a_. With 7.4×10^8^ (4.4×10^8^– 3.1×10^9^) s^-1^, outward-facing VGLUT1 releases glutamate faster than aspartate with 1.0×10^4^ (8.4×10^3^–1.4×10^4^) s^-1^ with similar binding rates (Figure 13 F, Figure 13— figure supplement 3).

To ensure coupled H^+^-glutamate transport, the glutamate translocation rate constant was limited to 1 s^-1^ for doubly protonated VGLUT1. Under these restrictions, translocation is much faster for aspartate with 8.5×10^4^ (6.9×10^4^–1.2×10^5^) s^-1^ than for glutamate with 1.4 (0.3–6.2) s^-1^ while singly protonated outward translocation rates are equivalent (Figure 13 D). This is consistent with the larger aspartate currents at very negative voltages in whole-cell recordings (Kolen, Borghans et al. 2023).

### Aspartate binding induces an occluded state that impairs Cl^-^ binding/unbinding

VGLUT1_PM_ binds Cl^-^ in different protonation and substrate-bound states (Figure 14, the corresponding *z* and *d* parameters are given in Figure 14—figure supplement 1). In transporters with no substrate bound, Cl^-^ binding has more pronounced effects on protonation of inward-facing states than of outward-facing states (Figure 14 A). For inward-facing conformations, Cl^-^ decreases the p*K*_a_ of the first protonation from 0.5 (0.4–0.7) to 6.3 (6.0–6.6), but increases the p*K*_a_ of the second protonation from 4.0 (3.8–4.0) to 7.3 (7.2–7.5). For outward-facing conformations, Cl^-^ slightly increases p*K*_a_ values for the first and second protonation states, from 7.8 (7.7–7.9) to 8.2 (8.1– 8.4) and from 7.7 (7.5–7.9) to 8.2 (8.1–8.4), respectively. Both the p*K*_a_ values and their Cl^-^ dependence are different from those of the protonation sites involved in VGLUT1 anion channel opening (Figure 4), consistent with the notion that different protonation sites are used for channel activation and transporter translocation.

**Figure 14:**
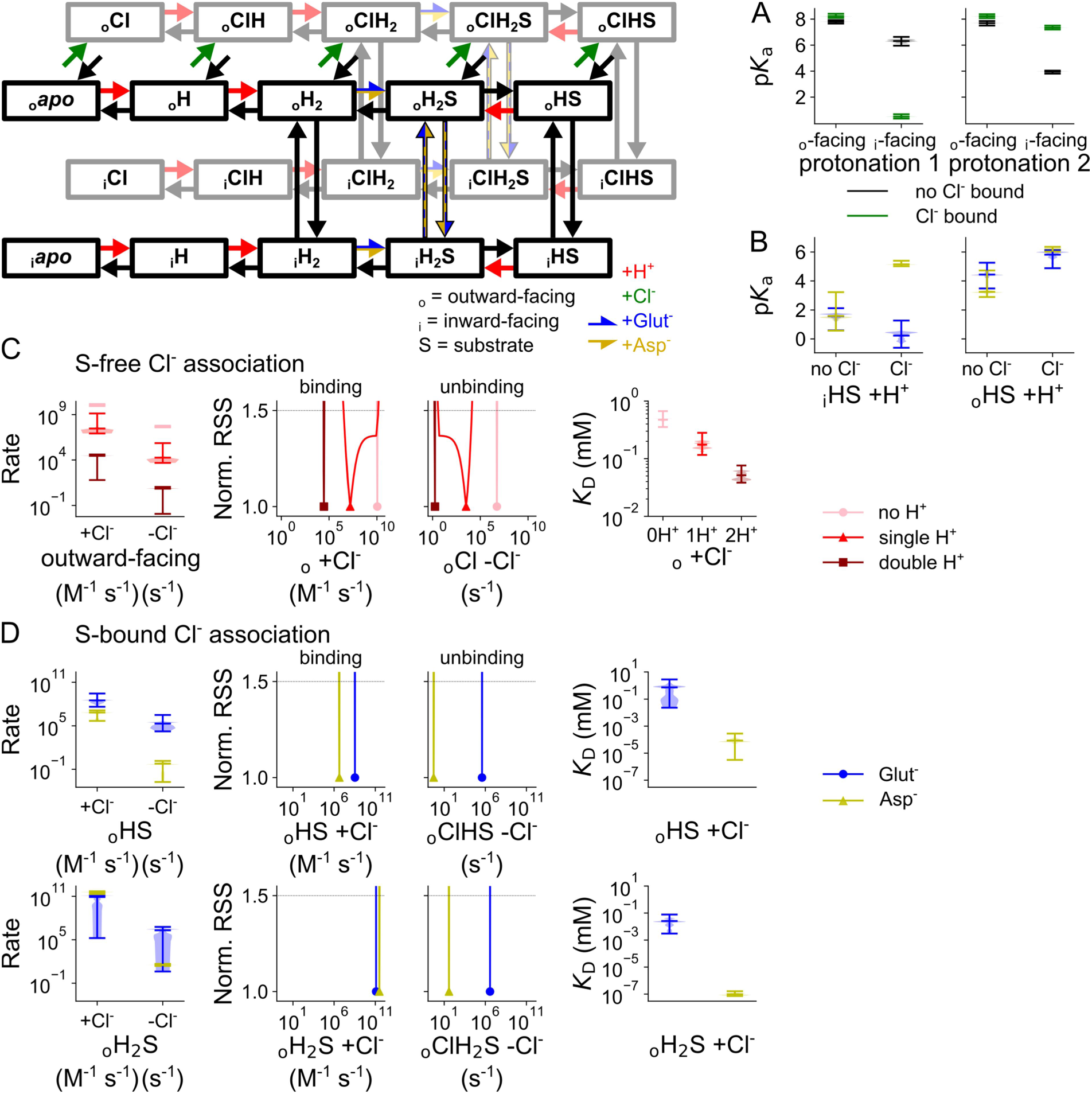
Binding and unbinding of protons and Cl^-^ to VGLUT1 in its neurotransmitter transport mode. p*K*_a_ for the first and second protonation, with and without Cl^-^, for inward- and outward-facing transporters without substrate (A), p*K*_a_ of the second protonation with bound substrate, with or without bound Cl^-^ (B), Cl^-^ binding/unbinding rates and corresponding *K*_D_ values for outward-facing transporters without substrate (light→dark red indicates increasing protonation; C), and Cl^-^ binding/unbinding rates and corresponding *K*_D_ values for the singly or doubly protonated, substrate-bound, outward-facing state (D). Protonation is represented by p*K*_a_ violin plots; other simulated rates are given as normalized RSS representing goodness of fit for a range of amplitudes in addition to violin plots depicting the amplitude range generated by exploratory mutation. All rates are shown at -160 mV.

**Figure 15:**
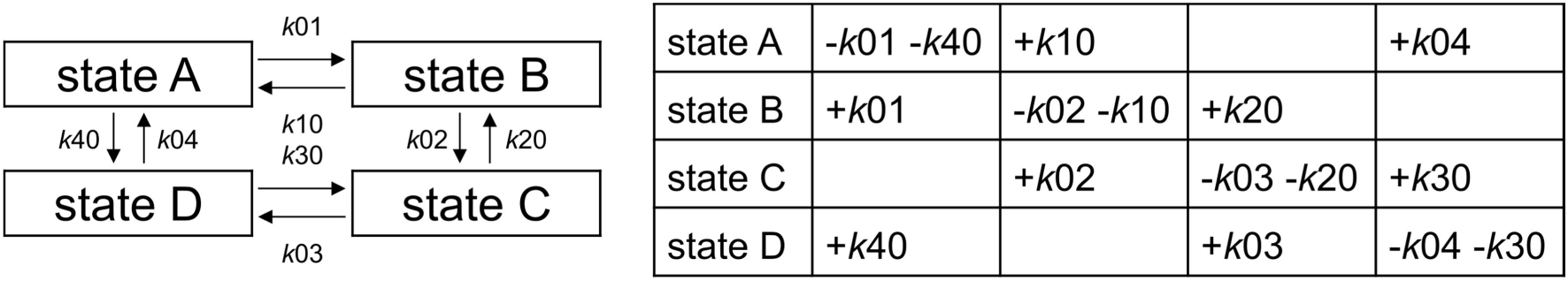
Hypothetical kinetic model and corresponding transition matrix. Directional rate numbers are chosen arbitrarily and full matrix equations are omitted for clarity. Any ligand binding rate would additionally be multiplied by the concentration of k0X, according to the convention used, for both its appearances in the matrix.

Cl^-^ binding differentially modifies the protonation of substrate-bound conformations (Figure 14 B): when glutamate is bound, Cl^-^ has no effect, but, when aspartate is bound, Cl^-^ increases the p*K*_a_ of the second protonation state for inward- and outward-facing transporters from 1.5 (0.6–3.2) to 5.1 (5.0–5.4) for _i_HS, and from 3.2 (2.9–4.7) to 6.2 (6.0–6.4) for _o_HS (Figure 14 B). Thus, Cl^-^ influences the substrate selectivity of VGLUTs by promoting protonation of the inward-facing aspartate-bound transporter such that aspartate mainly translocates with doubly protonated transporters in the uniport mode.

Cl^-^ unbinding from transporters without substrate is promoted by deprotonation, despite a simultaneous but smaller increase in binding. With 0.48 (0.35–0.67) mM Cl^-^ the non-protonated state has the highest *K*_D_, with a higher affinity of 0.18 (0.12–0.28) mM in the singly protonated state and 0.051 (0.038–0.076) mM in the doubly protonated state (Figure 14 C). However, statistical testing only identified significant differences in individual Cl^-^ association rates for substrate-bound transporters in the singly protonated state (_o_HS; Figure 14—figure supplement 2).

For substrate-bound VGLUT1 in the single protonation state, Cl^-^ binding and unbinding is much slower with bound aspartate than with bound glutamate: Cl^-^ binding with 1.1×10^4^ (7.7×10^2^–2.2×10^4^) M^-1^ s^-1^ versus 5.6×10^5^ (6.8×10^4^–4.6×10^6^) M^-1^ s^-1^ and Cl^-^ unbinding at 0.529 (0.002–1.433) s^-1^ versus 1.9×10^5^ (1.7×10^4^–3.1×10^6^) s^-1^ (Figure 14 D). These changes are directly observable in fast Cl^-^ application experiments (Figure 9 F), and indicate that binding of aspartate, but not of glutamate, promotes the formation of an occluded state that is partially inaccessible to Cl^-^ in the external solution. For double protonation, the Cl^-^ binding/unbinding rates for the glutamate- and aspartate-bound states are much closer; however, aspartate binding results in tighter Cl^-^ binding with a *K*_D_ of 9.5×10^-8^ (7.2×10^-8^–1.6×10^-7^) mM compared to glutamate with a *K*_D_ of 0.024 (0.003–0.078) mM (Figure 14 D).

## Discussion

Sustained synaptic activity requires the rapid recycling and refilling of synaptic vesicles. In glutamatergic synapses, VGLUTs fulfill two major refilling tasks: they accumulate glutamate and—to ensure that vesicular filling is electrically and osmotically neutral—mediate the efflux of Cl^-^, which is present at a high concentration in vesicles after endocytosis (Schenck, Wojcik et al. 2009, Eriksen, Chang et al. 2016, Martineau, Guzman et al. 2017, Kolen, Borghans et al. 2023). Both transport functions are regulated by luminal [Cl^-^] (Naito and Ueda 1985, Maycox, Deckwerth et al. 1988, Moriyama and Yamamoto 1995, Juge, Gray et al. 2010, Chang, Eriksen et al. 2018), and we studied the kinetic basis for this. We found that Cl^-^ accelerates H^+^-glutamate exchange, mainly by making the glutamate-binding site accessible to the cytoplasm, and by stimulating the inward translocation after substrate release to the vesicular lumen (Figure 12). Moreover, Cl^-^ increases the p*K*_a_ of the second protonation site of the aspartate-bound inward-facing transporter, thereby promoting double protonation and preventing H^+^-coupling of aspartate transport (Figure 14 B). During the activation of VGLUT1 anion channels, Cl^-^ stimulates the first protonation of the closed channel, accelerates opening of the singly protonated closed channel, and then stabilizes the open state by promoting protonation of the singly protonated open state and virtually abolishing channel closure regardless of protonation (Figure 4). The H120A point mutation is reported to modify VGLUT1 Cl^-^ channel gating and anion conduction (Kolen, Borghans et al. 2023). We found this mutation to increase the H^+^-binding affinity of the *apo* state well above to the levels observed in the WT, especially in the absence of Cl^-^, and to impair Cl^-^ binding by reducing its binding to the closed channel and increasing unbinding from the open channel for all protonation states (Figure 8). Therefore, our results identified the mechanistic basis of the allosteric control of VGLUT1 function by Cl^-^ and confirmed the functional importance of this regulatory mode.

We studied VGLUT1 anion channel activity in the absence of glutamate (Eriksen, Chang et al. 2016, Kolen, Borghans et al. 2023) and neurotransmitter transport after the complete substitution of intracellular anions with glutamate/aspartate at negative voltages with negligible Cl^-^ influx. These conditions enabled us to separately analyze VGLUT1 anion channel function and glutamate/aspartate transport. We developed distinct kinetic schemes to describe the two sets of experiments: anion channel function was described with a kinetic scheme that includes three VGLUT1 protonation states, each with a closed and open anion channel conformation and all of which can bind Cl^-^ (Figure 2); glutamate and aspartate transport was described with an alternating access transport scheme (Figure 10). It is likely that VGLUT1 anion channels can also open within the glutamate transport cycle under physiological conditions. However, generating a kinetic scheme for the combined transport/channel cycle will require better structural insight into the formation of anion channels than currently available.

Both VGLUT1 transport functions depend on H^+^ and Cl^-^ association/dissociation rates that are beyond the experimental time resolution, and there are protonation steps with p*K*_a_ values outside pH values tolerated in our experiments. Kinetic modeling permits extending these parameter ranges by fitting time courses and concentration dependences under various experimental conditions. For glutamate/aspartate transport as well as for anion channel function, we used the minimal models sufficient to describe the existing experimental data. Our kinetic models do not separately describe each proton binding/unbinding process or conformation transition but, instead, combine distinct processes for which experimental data do not permit separation. Since the gating of VGLUT1 anion channels needs to be described with at least two protonation states, we also assumed two H^+^-binding sites in the transporter mode. VGLUTs contain many titratable residues (Li, Eriksen et al. 2020), and more than two of these might be protonated during VGLUT transport/channel functions. Protonation sites in the channel and transporter modes indeed differed in p*K*_a_ values as well as in Cl^-^ modulation (Figure 6 and Figure 14), indicating that different sites determine the pH dependence of these two functions in VGLUTs.

Many titratable VGLUT residues project into water-filled vestibules (Li, Eriksen et al. 2020), and electrostatic interactions with Cl^-^ or with negatively charged amino acids might modify their p*K*_a_ values (Baran, Chimenti et al. 2008, Castañeda, Fitch et al. 2009, Isom, Castañeda et al. 2011). During the activation of VGLUT1 anion channels, Cl^-^ increases the p*K*_a_ values of the first protonation site in the closed, but not in the open, conformation and increases the p*K*_a_ values of the second protonated open, but not the corresponding closed conformation (Figure 4). These differences suggest that Cl^-^ binds near the first protonation site, when the VGLUT1 anion channel is in the closed conformation. For the second protonation step, Cl^-^ appears to bind at a nearby site in the open, but not the closed, state. Resolving the open channel conformation at the molecular level may help to identify the VGLUT protonation sites involved in anion channel activation. Cl^-^ has additional effects on the opening and closing rates of VGLUT1_PM_ anion channels, and similar effects of permeating ions have already been described for many ion channels. Cl^-^ binding may affect hydrophobic gating or prevent channel closing by a steric interaction via a foot-in-the-door mechanism (Swenson and Armstrong 1981, Machtens, Kortzak et al. 2015).

Kinetic modeling provided new mechanistic insight into VGLUT anion channel activation: at neutral luminal pH, none of the responsible protonation sites is protonated and opening rates are negligible. Furthermore, Cl^-^ association rates are close to the diffusion limit, consistent with the outward-facing conformation VGLUT2 in recent cryogenic electron microscopy (cryo-EM) findings (Li, Eriksen et al. 2020). Luminal Cl^-^ supports protonation of the first site in the closed conformation; however, this induces a conformational state with impaired Cl^-^ accessibility from the external medium (Figure 5). Thus, the predominant anion channel activation pathway includes Cl^-^ binding to unprotonated VGLUT1_PM_, followed by protonation, transition into an occluded state and, finally, channel opening. VGLUT1_PM_ anion channels are voltage-dependent, and channel opening upon membrane hyperpolarization and channel closure upon depolarization form the basis for the strong rectification of VGLUT1_PM_ Cl^-^ currents (Kolen, Borghans et al. 2023). Membrane hyperpolarization decreases the p*K*_a_ values for closed-state unprotonated channel by one or two units, but increases them for the second protonation (Figure 4 A). Negative voltages promotes opening rates of singly protonated channels by 2–3 orders of magnitude, but do not increase the opening rates of the doubly protonated channel. Taken together, the results show that membrane hyperpolarization increases the open probability of the VGLUT1_PM_ anion channel by promoting the opening rates of the singly protonated state as the primary opening pathway, despite also hampering the initial protonation.

To describe VGLUT1_PM_ glutamate/aspartate transport we used an alternating access scheme (Figure 10). VGLUT1_PM_ can transport many different anions, but glutamate transport is uniquely coupled to H^+^ exchange (Kolen, Borghans et al. 2023). To account for this behavior, we assumed that glutamate would be transported across the membrane in a distinct transporter protonation state compared with other substrates (Figure 10). Glutamate and aspartate can modify VGLUT1 protonation in a similar allosteric fashion to Cl^-^ (Figure 13 and Figure 14).

Inward-facing transporters with bound Cl^-^ and aspartate exhibit a p*K*_a_ of 5.1 (5.0–5.4), thus making double protonation likely; glutamate-bound transporters exhibit a p*K*_a_ of 0.33 (-0.61–1.3), which promotes H^+^ release from the second protonation site (Figure 14 B). Since retranslocation after substrate release occurs in a doubly protonated conformation, the preferential double protonation of inward-facing aspartate-bound states is sufficient to ensure aspartate uniport. Figure 13—figure supplement 2 illustrates the impact of this particular protonation on glutamate and aspartate transport rates. Increasing the p*K*_a_ without further optimizing other rates results in steep reduction of the predicted H^+^-glutamate exchange rates, and decreasing the p*K*_a_ towards glutamate-bound values similarly decreases aspartate transport.

Structural analysis of VGLUT2 suggests that glutamate is coordinated by the arginine residues R88 and R322 (Li, Eriksen et al. 2020). Aspartate is shorter and cannot bind both side chains simultaneously. These differences in the binding pocket might affect the protonation of nearby residues and may explain why glutamate promotes deprotonation of the substrate-bound inward-facing conformation. However, without limiting glutamate translocation in the doubly protonated state, our model fails to predict exclusive translocation for the singly protonated glutamate-bound conformation (Figure 13 G). Therefore, we conclude that additional mechanisms ensure stoichiometrically coupled exchange, possibly steric effects that prevent glutamate-bound translocation in the doubly protonated conformation. The much slower activation time course of aspartate currents by Cl^-^ (Figure 9) indicates that—during the aspartate transport cycle—Cl^-^ binding either promotes a conformational change or requires a conformational change other than that induced by Cl^-^ binding to the glutamate transporter. A likely scenario is that intermediate occluded states are more stable when aspartate is bound rather than glutamate.

Together with inorganic phosphate transporters, the lysosomal H^+^/sialic acid cotransporter sialin, and the vesicular nucleotide transporter VNUT, VGLUTs form the SLC17 family. Except for sialin (SLC17A5), all SLC17 transporters are regulated by Cl^-^ (Reimer 2013). Thus far, the physiological importance of this type of regulation remains insufficiently understood. Juge et al. (2010) demonstrated that ketone bodies block VGLUTs by competing with Cl^-^ at the allosteric regulation site and suggested that this regulatory mode will reduce glutamate release and inhibit network activity under conditions of insufficient food supply and increased ketone body production. Allosteric Cl^-^ regulation might also restrict glutamate accumulation in synaptic vesicles under physiological conditions. Neurons exhibit intracellular [glutamate] of 5– 10 mM (Danbolt 2001), and VGLUT1-mediated electrogenic H^+^-glutamate exchange may generate equilibrium glutamate concentrations of above 1 M in synaptic vesicles, causing water influx and vesicle swelling. The dual function as anion channel and transporter, together with allosteric Cl^-^ regulation, may prevent such glutamate overloading: VGLUT chloride channels deplete Cl^-^ from synaptic vesicles during glutamate filling, and the resulting low luminal [Cl^-^] will block glutamate uptake at a certain filling level.

H120A VGLUT1 has profoundly altered H^+^-glutamate coupling and anion channel gating (Kolen, Borghans et al. 2023), but kinetic modeling demonstrated only minor alterations in the kinetic scheme developed for WT VGLUT1 anion channels. The H120A mutation promotes closed channel protonation, especially without Cl^-^, but reduces protonation under most other conditions. The mutation reduces the opening rates of singly protonated channels, while increasing doubly protonated opening when Cl^-^ is bound. Moreover it increases closing rates, and these changes together reduce the stability of the singly protonated channel. This may explain why it takes longer to reach steady state current. In the open channel with no Cl^-^ bound, the H120A mutation causes no meaningful changes. These results demonstrate that

H120 is not one of the protonation sites necessary for anion channel opening. They rather indicate that Cl^-^ binding is impaired in virtually all protonation states of open and closed H120A VGLUT1_PM_ channels (Figure 8 C). Experimental (Chang, Eriksen et al. 2018) and computational (Rostamipour, Talandashti et al. 2022) work identified R176/R184 (VGLUT1/VGLUT2) as the luminal Cl^-^-binding site residue. In atomistic simulations, protonated H128 (in VGLUT2, corresponding to H120 in VGLUT1) was shown to support Cl^-^ binding to R184 (Rostamipour, Talandashti et al. 2022), in full support of our kinetic modeling results.

In summary, we have identified modification of H^+^ binding as a key process in VGLUT glutamate transport and anion channel activation. Transmembrane voltage promotes anion channel opening with single protonation, and luminal Cl^-^ increases the p*K*_a_ values to reach this protonation state as well as stabilizing open channels in general (Figure 4). Differences in transporter protonation contribute to the distinct transport stoichiometries of glutamate or aspartate: glutamate binding promotes H^+^ release to the cytoplasm and permits preferential glutamate translocation in the singly protonated state and stoichiometrically coupled H^+^-glutamate exchange, whereas aspartate binding does not (Figure 13 C). We found that Cl^-^ promotes conformational changes, most importantly by facilitating substrate binding from the cytoplasm (Figure 13—figure supplement 3) or by increasing anion channel opening and substantially decreasing channel closing rates (Figure 8 B). Analyzing the rates of Cl^-^ binding/unbinding demonstrated that Cl^-^ association is impaired in certain protonation states during channel activation (Figure 5 A), likely via the formation of occluded states or via translocation into inward-facing conformations. Taken together, our results illustrate how vesicular glutamate transporters use allosteric interactions between the anionic substrate glutamate, the main physiological anion Cl^-^, and the driving co-substrate H^+^ to orchestrate multiple functions of these key players in synaptic transmission.

## Materials and methods

### Expression plasmids, mutagenesis, and heterologous expression

HEK293T cells (Sigma-Aldrich) were grown in humidified incubators at 37°C and 95% air/5% CO_2_ in Dulbecco’s modified Eagle Medium supplemented with 10% FBS, and 5 mL penicillin-streptavidin at 5,000 U/mL. To express VGLUT1 as an eGFP fusion protein, we inserted the coding region of eGFP into pcDNA3.1-rVGLUT1 (kindly provided by Dr. Shigeo Takamori at Doshisha University in Kyoto, Japan) at the 3ʹ end of the transporter coding region and mutated various dileucine-like endocytosis motifs to alanine to promote plasma membrane insertion using PCR-based techniques (Kolen, Borghans et al. 2023). The H120A mutation was introduced into VGLUT1_PM_ using PCR-based techniques. Cells were transiently transfected using a calcium phosphate or lipofectamine precipitation method and examined 24 h later.

### Whole-cell patch clamp and fluorescence measurements

Standard whole-cell patch clamp recordings were performed using EPC10 amplifiers, controlled by HEKA PatchMaster (Multi Channel Systems MCS GmbH, Reutlingen, Germany; Kolen, Borghans et al. 2023). We used borosilicate pipettes (Harvard Apparatus, Holliston, Massachusetts, USA) with a resistance of 0.9–3 MΩ and applied series resistance compensation and capacitance cancelation, resulting in a voltage error of < 5 mV. The standard bath solution contained (in mM) 136 choline chloride, 2 MgCl_2_, and 30 HEPES or 50 MES; we adjusted the osmolarity of the bath solution with glucose to values at least 5 mOsm/L higher than the internal solution. HEPES was used as the buffer for external and internal pH values between 6.5 and 7.4, and was replaced with MES for more acidic perfusion solutions; in all cases, pH was adjusted with choline hydroxide. In experiments that used glutamate or aspartate in the pipette solution, 100 mM choline chloride was replaced by choline gluconate in the bath solution; cells were held at -50 mV before all measures other than the Cl^-^ current. The standard pipette solution contained (in mM) 140 choline anion_X_, 5 EGTA, 5 Mg(OH)_2_, and 30 HEPES, adjusted to pH 7.4 with TMA-OH (anion_X_ = Cl^-^, glutamate^-^ or aspartate^-^). For glutamate- or aspartate-based pipette solutions, we used internal agar salt bridges (made from plastic tubing filled with 0.5 M KCl in 2% agar) to connect the Ag/AgCl electrode. Where necessary, junction potentials were calculated and corrected. To support complete intracellular dialysis, we waited at least 2 min after establishing the whole-cell mode before starting electrical recordings. Except for Cl^-^ dependence of the anion channel function, current recordings were corrected for leakage currents by subtracting current amplitudes obtained in the same cell with external pH 7.4 under identical conditions. In an earlier publication (Kolen, Borghans et al. 2023), we carefully tested whether glutamate and aspartate currents measured in this way are mediated by VGLUT1_PM_: both are blocked by Rose Bengal (a high-affinity, non-competitive VGLUT blocker), only negligible currents can be recorded under these conditions in untransfected HEK293T cells, and there is a linear relationship between expression levels and transport currents.

Fast solution exchanges were achieved by moving the solution interface of continuous perfusion flow around a cell using a PiezoMove P-601.30L or P-840.40 piezoelectric actuator (Physik Instrumente, Karlsruhe, Germany) controlled via a MXPZT controller (Siskiyou, Grants Pass, Oregon, USA) or PI E-836 Piezo Driver combined with a USBPGF-S1 Instrumentation Amplifier Low Pass Filter (Alligator Technologies, Charlottesville, Virginia, USA) to smoothen the signal and reduce vibration. Actuators were connected to the EPC10 amplifier and controlled using HEKA PatchMaster software. The mean time course of solution exchange was determined to be 0.87 ± 0.2 ms (n = 19, mean and 95% confidence interval) by recording the current responses of an open pipette after changing from a high to low [Cl^-^] solution. Two back-and-forth exchanges were performed, and the on-off pair with the best resolution was chosen and averaged.

### Kinetic modeling

VGLUT chloride or glutamate/aspartate currents were simulated using kinetic models based on subsequent conformational changes and ligand binding/reactions. Changes in state occupation were computed via a matrix of rate equations that describe interactions between two connected states.

Solving this matrix as a series of linear algebra equations provided steady-state distributions of kinetic states. The dependence of net rates on voltage and temperature, as an adaptation of the Arrhenius equation, is calculated as

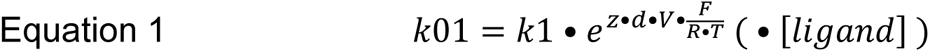

where *k*1 is the rate constant (limited to < 10^5^ s^-1^ for conformation changes and to 5×10^9^ for binding and unbinding of ligands), *z* is charge movement (-1 to +1), *d* is a symmetry factor (0 to 1, equivalent to *α* or *β* in the Butler–Volmer equation (Björketun et al. 2012), V is the voltage across the cell membrane, and the other elements are constants: *F* is the Faraday constant, *R* is the gas constant, and *T* is the absolute temperature. The inverse reaction *k10* was calculated as:

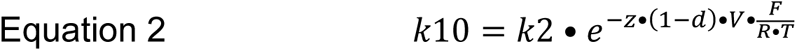

To ensure microscopic reversibility, one rate constant was calculated from the clockwise and counterclockwise products, and one *z* variable was the sum of all other *z* values in each cycle. To prevent transporter behavior that is incompatible with experimental data, the rate constants describing unprotonated channel opening and doubly protonated glutamate-bound transporter transition between inward- and outward-facing conformation, each with or without bound Cl^-^, were limited to 1 s^-1^. The calculated net rates may exceed 1 s^-1^ at -160 mV depending on their voltage dependence.

For current predictions in the VGLUT1_PM_ channel mode, the distribution of relative state occupations in the transition matrix was used to calculate the fraction of open states, i.e. the open probability, as the output value. Numerical weights were assigned to individual simulated metrics and time courses to control the driving force optimizing simulation accuracy through parameter mutation. A large weight ensures that the WT model remains close to the calculated open probability of 0.24 at 140 mM Cl^-^, pH 5.5, and -160 mV (Kolen, Borghans et al. 2023). Each simulation calculates the modification of steady-state currents by [Cl^-^]_o_, pH, or voltage relative to this open probability and the time-dependent changes of currents relative to the highest steady-state current in each experiment. Active transport was calculated as the sum of net transition fluxes between states, multiplied by the corresponding *z* value. Here, a microscopic reversibility charge offset of -1 or -2 in each transport cycle accounts for current generation. Similarly to how Cl^-^ channel models were trained to mimic experimentally determined open probability, the active transport model uses the calculated glutamate (561 s^-1^) and aspartate (2581 s^-1^) transport rates. Parameters were optimized via a population-based optimization procedure of the genetic algorithm of the Distributed Evolutionary Algorithms in Python (DEAP) software package (Fortin et al., 2012).

We converted the kinetic transition rates into transition probabilities in order to allow for the calculation of preferred pathways of opening and closing of the anion channel (Figure 3 L and Figure 7 L), using the reactive flux module of the Deeptime Python library which uses transition path theory (Metzner, Schütte et al. 2009). Activation pathways were described starting from the *apo* state to any conductive state, deactivation pathways from the two most common open states (oH or oClH_2_) to either of the two unprotonated closed states (*apo* and cCl).

### Quantification and statistical analysis

Data are given either as the mean ± 95% confidence interval or as violin plots showing the distribution of data with the median and exploratory mutation value range. We calculated secondary binding rates for Cl^-^, glutamate, and aspartate to normalize them, dividing the binding rate constant by the ligand concentration: in mM when calculating *K*_D_ (expressed in mM) and in M for the rates themselves (expressed in M^-1^ s^-1^).

Measurements were taken from distinct samples. Mean and confidence intervals of time courses, dependence fits, and fit parameters were either from individual fits (chloride dependence of glutamate and aspartate) or determined by bootstrapping (other dependence fits and time constants). For patch clamp electrophysiology experiments, sample sizes correspond to the number of patched cells and are given in the figure or legend. For statistical analysis of two groups, we used two-tailed bootstrap hypothesis testing via the bootstrap t-statistic of bootstrapped means, F-tests to determine the number of exponents in current activation and deactivation, and one-sample Wilcoxon signed rank tests when comparing fits of means to distributions of groups of individually fitted cells.

We fitted time courses of VGLUT1_PM_ currents with mono- or biexponential functions and concentration dependence by global bootstrapping, with the Michaelis– Menten equation for Cl^-^ and the Hill equation for pH to better capture saturation. To generate modest and comparable confidence intervals, we used sampling of 1000 for dependence data and time constants. To ensure reliable p-values, we used higher sampling of 10,000 for bootstrap hypothesis testing between different datasets.

For comparing simulated values and rates we used two tests, and changes in kinetic scheme parameters were only considered significant after passing both tests. A modified DEAP cycle version was used to collect all parameter set variants with RSS values within 125% for all simulated metrics and time courses. This exploratory mutation applied to the Cl^-^ current model generated >10,000 sets with >3000 variants for each parameter. In case of the glutamate/aspartate transport, we limited the data size by removing individuals that did not vary parameters describing protonation, and generated >980 variants from >20,000 individual sets from >50,000 generations.

All unique parameters values are shown in violin plots and are represented by the median and the range of amplitude variation generated during exploratory mutation as a confidence interval, in the text as well as in figures. Moreover, we calculated the relative RSS for a range of values centered on the optimized value for each individual parameter. If the values overlapped at RSS of below 150%, the parameters were assumed to be indistinct.

## Code Availability

Kinetic models, parameters, the simulation source code as well as instructions are available at https://jugit.fz-juelich.de/computational-neurophysiology/vglut-kinetic-models

## Acknowledgments

We thank Dr. Shigeo Takamori for kindly providing pcDNA3.1-rVGLUT1 and Dr. B. Kolen for doing some of the measurements. The authors gratefully acknowledge the computing time on the supercomputer JURECA at Forschungszentrum Jülich granted through JARA under grant no. mpogt. This work was supported by the Deutsche Forschungsgemeinschaft (German Research Foundation) to Ch.F. (FA 301/15–2 as part of Research Unit FOR 2518 *DynIon* and FA 301/13-2 as part of the Research Unit FOR 2795 *Synapses under stress*).

## Author Contributions

B.B. designed and performed most of the experiments and kinetic modeling; D.K. contributed to the kinetic modeling; P.L. and J.-P.M. supported optimization of the model and simulation code for high-performance computing; Ch.F. conceived the idea, designed the experiments, supervised the work, and wrote the manuscript.

## Competing interests

The authors have no conflicts of interest.

## Notes

https://jugit.fz-juelich.de/computational-neurophysiology/vglut-kinetic-models

### Competing Interest Statement

The authors have declared no competing interest.

### Summary of Updates

Text provided as PDF to avoid the platform's converter from degrading images.

## References

Baran, K. L., M. S. Chimenti, J. L. Schlessman, C. A. Fitch, K. J. Herbst and B. E. Garcia-Moreno (2008). “Electrostatic effects in a network of polar and ionizable groups in staphylococcal nuclease.” J Mol Biol 379(5): 1045–1062.

Bergles, D. E., A. V. Tzingounis and C. E. Jahr (2002). “Comparison of coupled and uncoupled currents during glutamate uptake by GLT-1 transporters.” Journal of Neuroscience 22(23): 10153–10162.

Bhat, S., M. Niello, K. Schicker, C. Pifl, H. H. Sitte, M. Freissmuth and W. Sandtner (2021). “Handling of intracellular K^+^ determines voltage dependence of plasmalemmal monoamine transporter function.” Elife 10; e67996.

Castañeda, C. A., C. A. Fitch, A. Majumdar, V. Khangulov, J. L. Schlessman and B. E. García-Moreno (2009). “Molecular determinants of the pKa values of Asp and Glu residues in staphylococcal nuclease.” Proteins 77(3): 570–588.

Chang, R., J. Eriksen and R. H. Edwards (2018). “The dual role of chloride in synaptic vesicle glutamate transport.” Elife 7.

Danbolt, N. C. (2001). “Glutamate uptake.” Progress in Neurobiology 65(1): 1–105.

Eriksen, J., R. Chang, M. McGregor, K. Silm, T. Suzuki and R. H. Edwards (2016). “Protons regulate vesicular glutamate transporters through an allosteric mechanism.” Neuron 90(4): 768–780.

Ewers, D., T. Becher, J. P. Machtens, I. Weyand and C. Fahlke (2013). “Induced fit substrate binding to an archeal glutamate transporter homologue.” Proc Natl Acad Sci U S A 110(30): 12486–12491.

Farsi, Z., R. Jahn and A. Woehler (2017). “Proton electrochemical gradient: Driving and regulating neurotransmitter uptake.” Bioessays 39(5): doi: 10.1002/bies.201600240

Franke, C., H. Hatt and J. Dudel (1987). “Liquid filament switch for ultra-fast exchanges of solutions at excised patches of synaptic membrane of crayfish muscle.” Neurosci Lett. 77(2): 199–204.

Grewer, C., N. Watzke, M. Wiessner and T. Rauen (2000). “Glutamate translocation of the neuronal glutamate transporter EAAC1 occurs within milliseconds.” Proceedings of the National Academy of Sciences of the United States of America 97(17): 9706–9711.

Hanke, W. and C. Miller (1983). “Single chloride channels from *Torpedo* electroplax. Activation by protons.” J Gen Physiol 82: 25–45.

Isom, D. G., C. A. Castañeda, B. R. Cannon and B. García-Moreno (2011). “Large shifts in pKa values of lysine residues buried inside a protein.” Proc Natl Acad Sci U S A 108(13): 5260–5265.

Jonas, P. and B. Sakmann (1992). “Glutamate receptor channels in isolated patches from CA1 and CA3 pyramidal cells of rat hippocampal slices.” J Physiol 455: 143–171.

Juge, N., J. A. Gray, H. Omote, T. Miyaji, T. Inoue, C. Hara, H. Uneyama, R. H. Edwards, R. A. Nicoll and Y. Moriyama (2010). “Metabolic control of vesicular glutamate transport and release.” Neuron 68(1): 99–112.

Kolen, B., B. Borghans, D. Kortzak, V. Lugo, C. Hannack, R. E. Guzman, G. Ullah and C. Fahlke (2023). “Vesicular glutamate transporters are H^+^-anion exchangers that operate at variable stoichiometry.” Nat Commun 14(1): 2723.

Kortzak, D., C. Alleva, I. Weyand, D. Ewers, M. I. Zimmermann, A. Franzen, J. P. Machtens and C. Fahlke (2019). “Allosteric gate modulation confers K^+^ coupling in glutamate transporters.” Embo j: e101468.

Kovermann, P., M. Hessel, D. Kortzak, J. C. Jen, J. Koch, C. Fahlke and T. Freilinger (2017). “Impaired K+ binding to glial glutamate transporter EAAT1 in migraine.” Sci Rep 7(1): 13913.

Lau, A. Y., H. Salazar, L. Blachowicz, V. Ghisi, A. J. Plested and B. Roux (2013). “A conformational intermediate in glutamate receptor activation.” Neuron 79(3): 492–503.

Li, F., J. Eriksen, J. Finer-Moore, R. Chang, P. Nguyen, A. Bowen, A. Myasnikov, Z. Yu, D. Bulkley, Y. Cheng, R. H. Edwards and R. M. Stroud (2020). “Ion transport and regulation in a synaptic vesicle glutamate transporter.” Science 368(6493): 893–897.

Machtens, J. P., D. Kortzak, C. Lansche, A. Leinenweber, P. Kilian, B. Begemann, U. Zachariae, D. Ewers, B. L. de Groot, R. Briones and C. Fahlke (2015). “Mechanisms of anion conduction by coupled glutamate transporters.” Cell 160(3): 542–553.

Martineau, M., R. E. Guzman, C. Fahlke and J. Klingauf (2017). “VGLUT1 functions as a glutamate/proton exchanger with chloride channel activity in hippocampal glutamatergic synapses.” Nat Commun 8(1): 2279.

Maycox, P. R., T. Deckwerth, J. W. Hell and R. Jahn (1988). “Glutamate uptake by brain synaptic vesicles. Energy dependence of transport and functional reconstitution in proteoliposomes.” J Biol Chem 263(30): 15423–15428.

Metzner, P., C. Schütte and E. Vanden-Eijnden (2009). “Transition Path Theory for Markov Jump Processes.” Multiscale Modeling & Simulation 7(3): 1192–1219.

Moriyama, Y. and A. Yamamoto (1995). “Vesicular L-glutamate transporter in microvesicles from bovine pineal glands. Driving force, mechanism of chloride anion activation, and substrate specificity.” J Biol Chem 270(38): 22314–22320.

Naito, S. and T. Ueda (1985). “Characterization of glutamate uptake into synaptic vesicles.” J Neurochem 44(1): 99–109.

Oh, S. and O. Boudker (2018). “Kinetic mechanism of coupled binding in sodium-aspartate symporter GltPh.” Elife 7.

Omote, H., T. Miyaji, M. Hiasa, N. Juge and Y. Moriyama (2016). “Structure, Function, and Drug Interactions of Neurotransmitter Transporters in the Postgenomic Era.” Annu Rev Pharmacol Toxicol 56: 385–402.

Otis, T. S. and M. P. Kavanaugh (2000). “Isolation of current components and partial reaction cycles in the glial glutamate transporter EAAT2.” Journal of Neuroscience 20(8): 2749–2757.

Reimer, R. J. (2013). “SLC17: a functionally diverse family of organic anion transporters.” Mol Aspects Med 34(2-3): 350–359.

Rostamipour, K., R. Talandashti and F. Mehrnejad (2022). “Atomistic insight into the luminal allosteric regulation of vesicular glutamate transporter 2 by chloride and protons: An all-atom molecular dynamics simulation study.” Proteins 90(12): 2045–2057.

Schenck, S., S. M. Wojcik, N. Brose and S. Takamori (2009). “A chloride conductance in VGLUT1 underlies maximal glutamate loading into synaptic vesicles.” Nat Neurosci 12(2): 156–162.

Schicker, K., Z. Uzelac, J. Gesmonde, S. Bulling, T. Stockner, M. Freissmuth, S. Boehm, G. Rudnick, H. H. Sitte and W. Sandtner (2012). “Unifying concept of serotonin transporter-associated currents.” J Biol Chem 287(1): 438–445.

Suslova, M., D. Kortzak, J. P. Machtens, P. Kovermann and C. Fahlke (2023). “Apo state pore opening as functional basis of increased EAAT anion channel activity in episodic ataxia 6.” Front Physiol 14: 1147216.

Swenson, R. P., Jr and C. M. Armstrong (1981). “K^+^ channels close more slowly in the presence of external K+ and Rb+.” Nature 291(5814): 427–429.

Takamori, S. (2016). “Vesicular glutamate transporters as anion channels?” Pflugers Arch 468(3): 513–518.

Wang, F., S. Redding, I. J. Finkelstein, J. Gorman, D. R. Reichman and E. C. Greene (2013). “The promoter-search mechanism of Escherichia coli RNA polymerase is dominated by three-dimensional diffusion.” Nat Struct Mol Biol 20(2): 174–181.

Zhang, Z., Z. Tao, A. Gameiro, S. Barcelona, S. Braams, T. Rauen and C. Grewer (2007). “Transport direction determines the kinetics of substrate transport by the glutamate transporter EAAC1.” Proc.Natl.Acad.Sci.U.S.A 104(46): 18025–18030.

