## supplemental figures for "Allosteric modulation of proton binding confers Cl^-^ activation and glutamate selectivity to vesicular glutamate transporters"

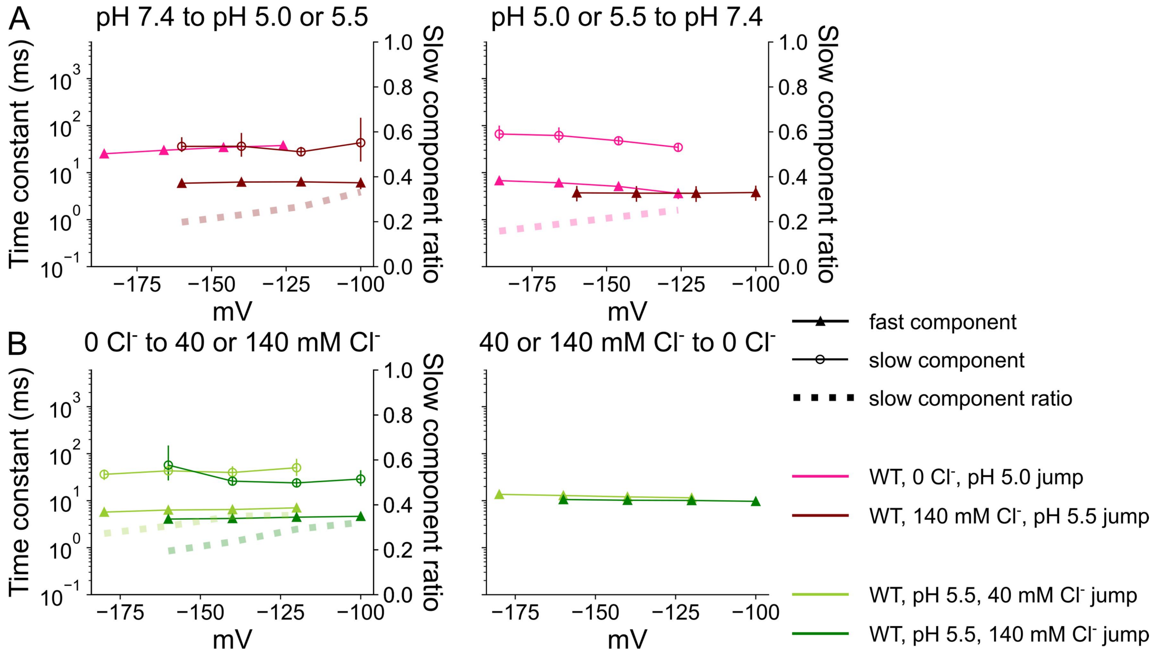


**Figure 1—figure supplement 1: Time constants for VGLUT1$_{PM}$ $Cl^-$ current activation/deactivation in response to $H^+$ or $Cl^-$ concentration steps.** Activation/deactivation time constants upon pH jumps from 7.4 to 5.0 or 5.5 (left) or from 5.0 or 5.5 to 7.4 (right) at an external [$Cl^-$] of 0 or 140 mM (**A**), or upon [$Cl^-$] steps from 0 to 40 mM or 0 to 140 mM at pH 5.5 (**B**). Activation at high external $Cl^-$ or deactivation without $Cl^-$ were fitted with biexponential functions, providing two time constants and the relative amplitude (dashed lines) of the slower component. Data are shown as means obtained by bootstrapping with a global fit of experimental data with a sampling of 1000, with 95% of the sampling as error bars. Voltage differences are the result of *a posteriori* liquid junction potential correction.

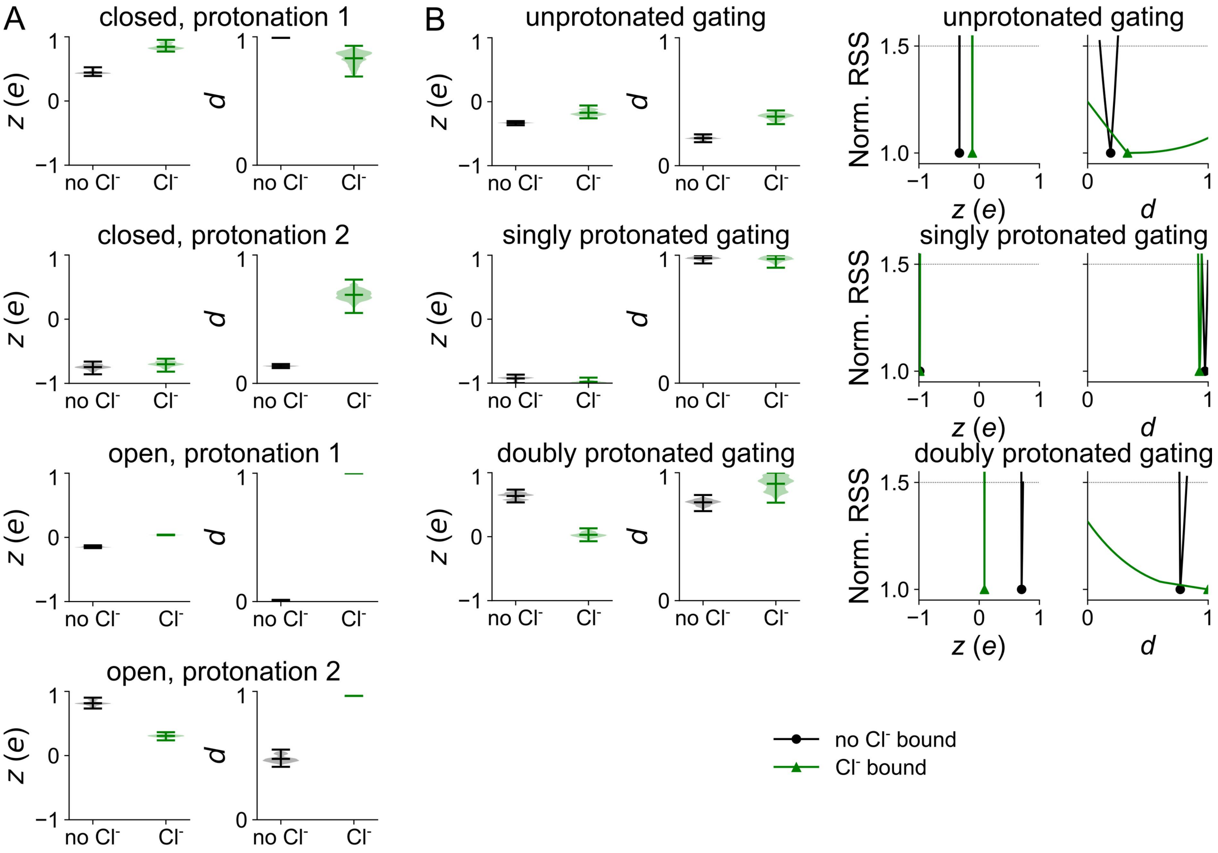


**Figure 4—figure supplement 1: Voltage dependence of parameters of the WT $VGLUT1_{PM}$ anion channel kinetic scheme.** Distribution of *z* and *d* parameters for protonation steps with and without $Cl^-$ (**A**), distribution of *z* and *d* parameters for channel opening (**B**). Protonation parameters are represented by violin plots, other simulation results are given as normalized RSS representing goodness of fit for a range of amplitudes in addition to violin plots depicting the amplitude range generated by exploratory mutation.

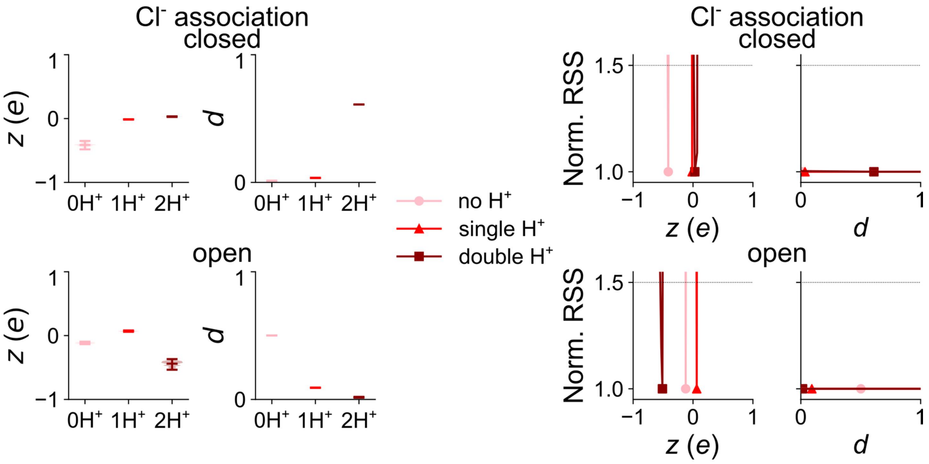


**Figure 5—figure supplement 1: Voltage dependence of Cl$^-$ association parameters of the WT VGLUT1$_{PM}$ anion channel kinetic scheme.** Distribution of $z$ and $d$ parameters for the three protonation states (light→dark red indicates increasing protonation). Simulation results are given as violin plots depicting the amplitude range generated by exploratory mutation and normalized RSS representing goodness of fit for a range of amplitudes.

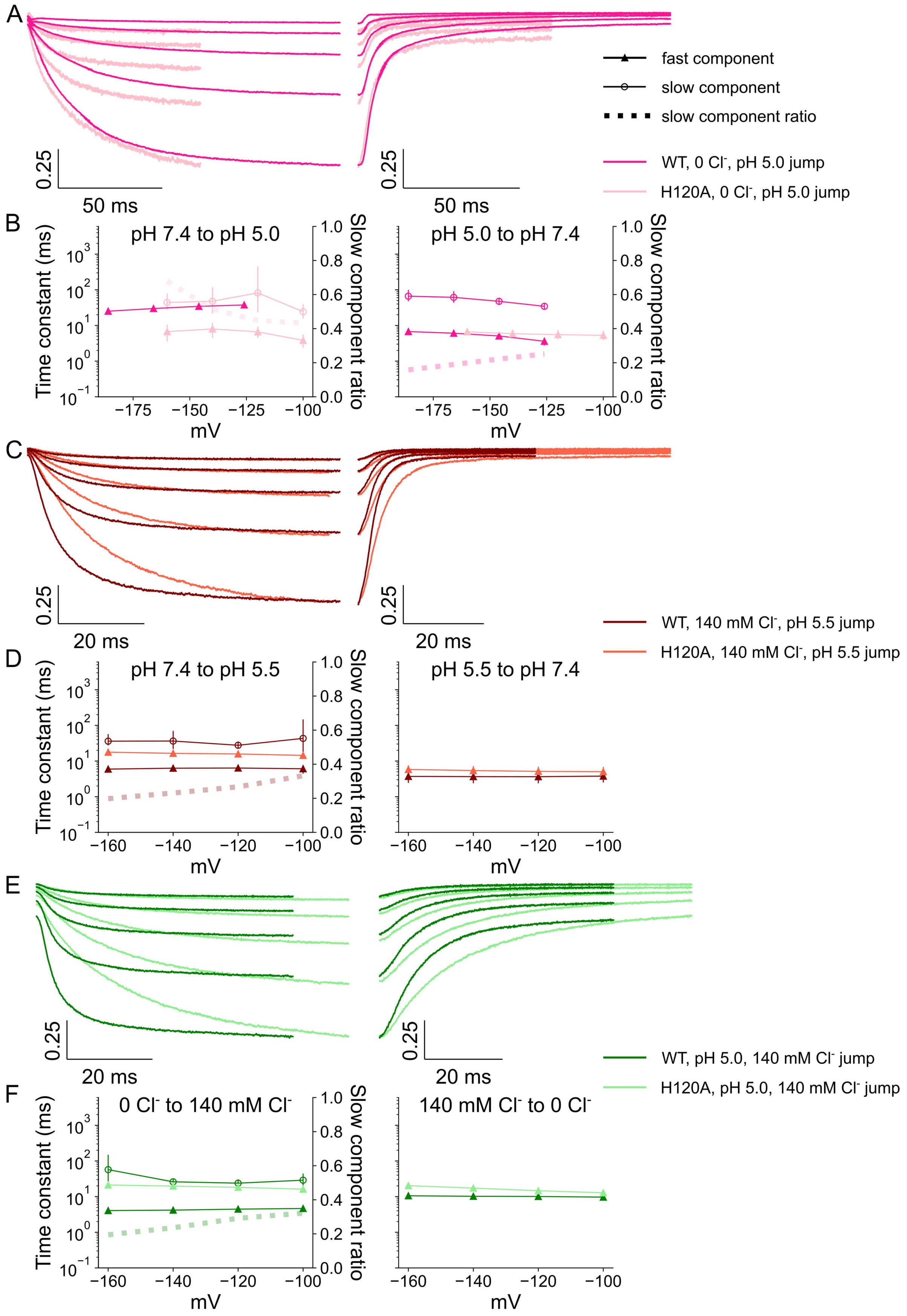


**Figure 6—figure supplement 1: Comparison of WT and H120A VGLUT1 anion current kinetics.** Representative current responses to concentration jumps from pH 7.4 to 5.0 at [Cl$^-$] = 0 mM (**A**) with corresponding time constants for activation (left)

and deactivation (right; **B**), representative current responses to concentration jumps from pH 7.4 to 5.0 at [$Cl^-$] = 0 mM140 mM (**C**) with corresponding time constants for activation (left) and deactivation (right; **D**), and representative current responses to concentration jumps from [$Cl^-$] = 0 to 140 mM at pH 5.5 (**E**) with corresponding time constants for activation (left) and deactivation (right; **F**) at given holding potentials. Activation at high external $Cl^-$ or deactivation without $Cl^-$ were fitted with biexponential functions, with two time constants and the relative amplitudes (dashed lines) of the slower component shown. Data are shown as means obtained by bootstrapping with a global fit of experimental data with a sampling of 1000, with 95% of the sampling as error bars. Voltage differences are the result of *a posteriori* liquid junction potential correction.

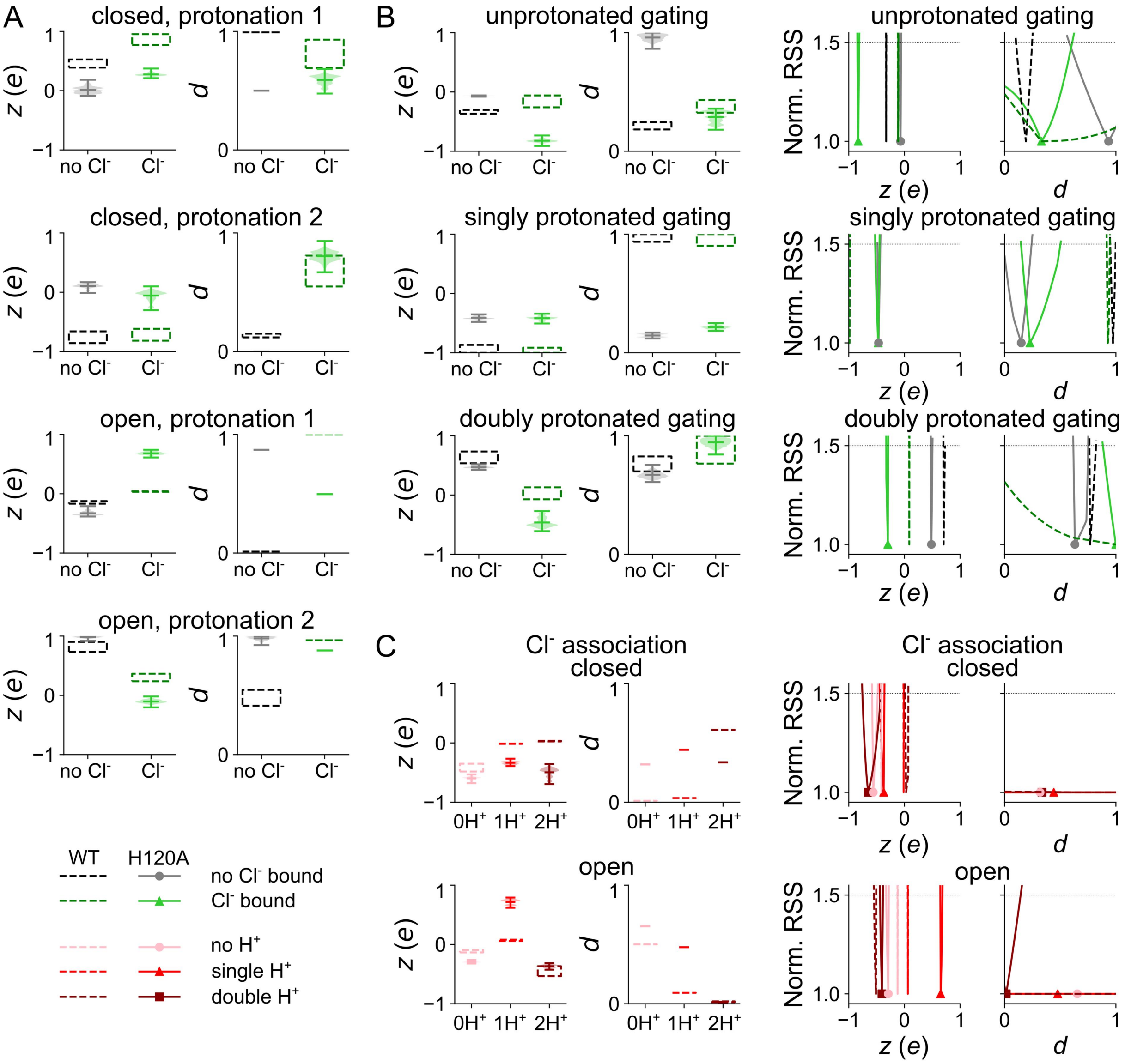


**Figure 8—figure supplement 1: Voltage dependence of kinetic parameters of the H120A VGLUT1$_{PM}$ anion channel kinetic scheme.** Distribution of *z* and *d* parameters for protonation steps with and without $Cl^-$ (**A**), for channel opening with and without $Cl^-$ (**B**), and for $Cl^-$ binding by protonation state (**C**). Protonation parameters are represented by violin plots, other simulation results are given as normalized RSS representing goodness of fit for a range of amplitudes in addition to violin plots depicting the amplitude range generated by exploratory mutation.

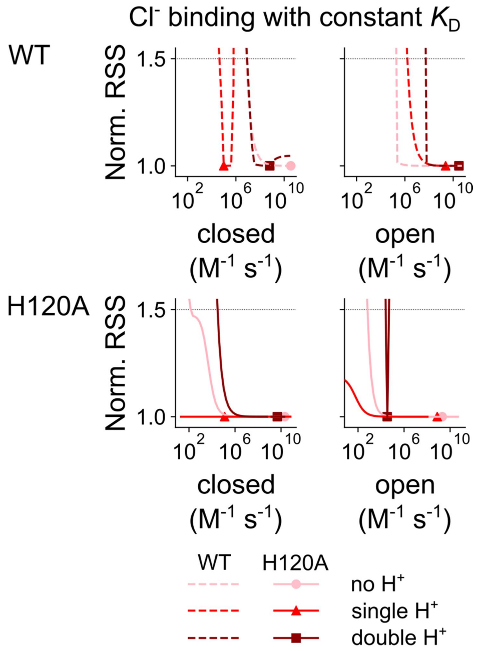


**Figure 8—figure supplement 2: Statistical analysis of $Cl^-$ binding by WT and H120A VGLUT1$_{PM}$ at a constant $K_D$.** Changes in the goodness of fit upon the modification of secondary $Cl^-$-binding rate constants at -160 mV, to closed (left) or open (right) WT or H120A VGLUT1$_{PM}$ anion channels in the unprotonated or singly or doubly protonated state. During modification, unbinding constants were simultaneously altered to keep the $Cl^-$-binding affinity unchanged.

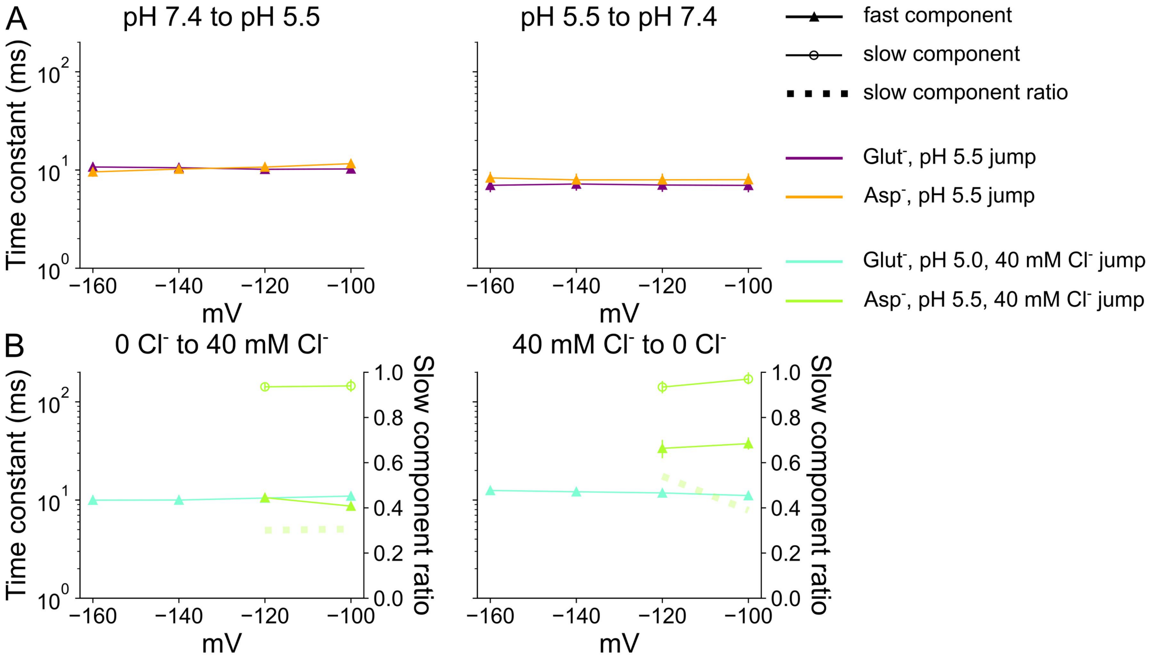


**Figure 9—figure supplement 1: Time constants for VGLUT1$_{PM}$ glutamate or aspartate current activation/deactivation by $H^+$ or $Cl^-$.** Activation/deactivation time constants upon pH jumps from 7.4 to 5.0 (left) or deactivation time constants upon pH jumps from 5.0 to 7.4 (right) at an external [Cl$^-$] of 40 mM (**A**) or upon [Cl$^-$] jumps from 0 to 40 mM (left) or deactivation time constants upon [Cl$^-$] jumps from 40 to 0 mM (right) at an external pH of 5.0 for glutamate or 5.5 for aspartate (**B**). Activation and deactivation of aspartate currents by $Cl^-$ were fitted with biexponential functions, providing two time constants and relative amplitudes (dashed lines) for the slower component. Data are shown as means obtained by bootstrapping with a global fit of experimental data with a sampling of 1000, with 95% of the sampling as error bars.

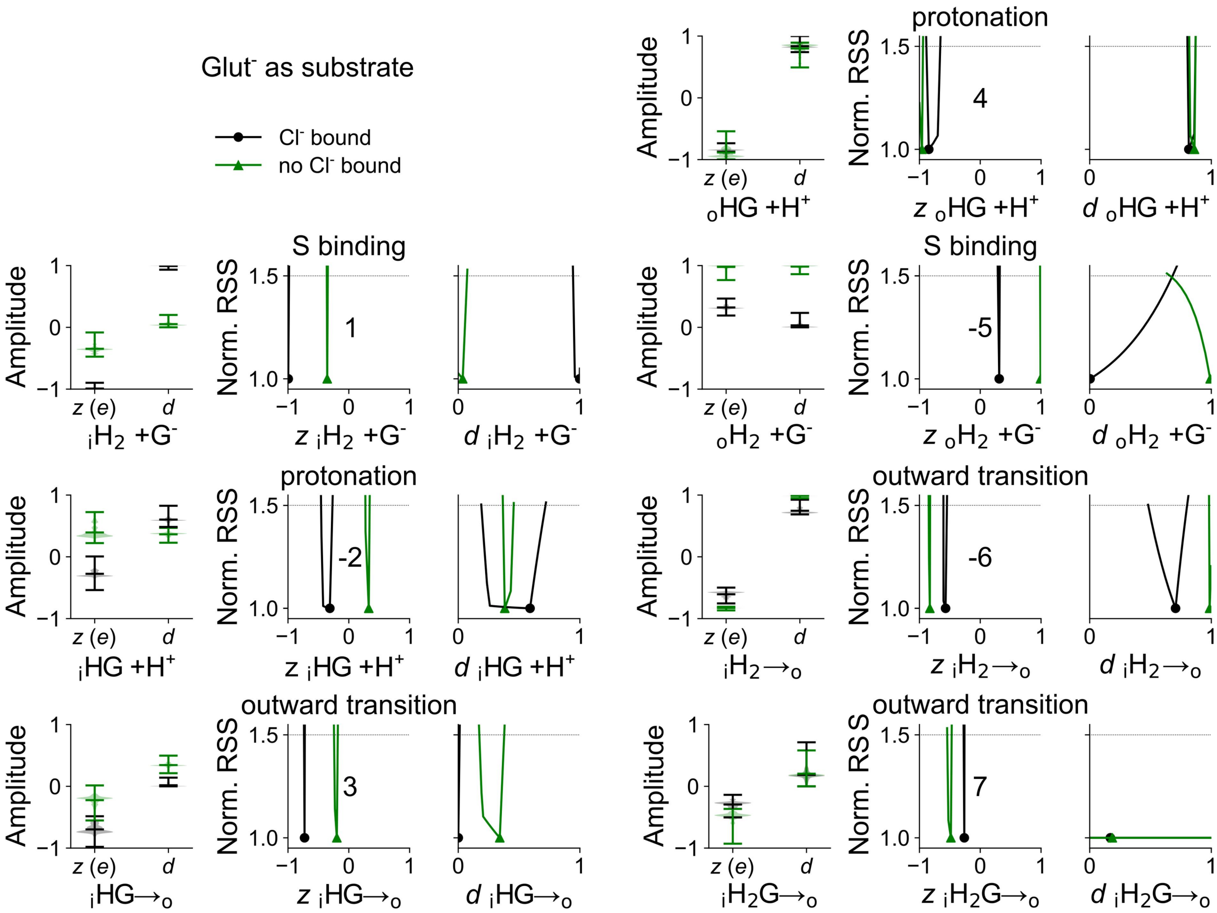


**Figure 12—figure supplement 1: Voltage dependence of parameters of the glutamate transport cycles.** Distribution of *z* and *d* parameters for steps describing substrate binding (1 and 5), protonation (2 and 4), and transition between inward- and outward-facing conformations (3, 6, and 7). Simulation results are given as violin plots depicting the amplitude range generated by exploratory mutation and normalized RSS representing goodness of fit for a range of amplitudes.

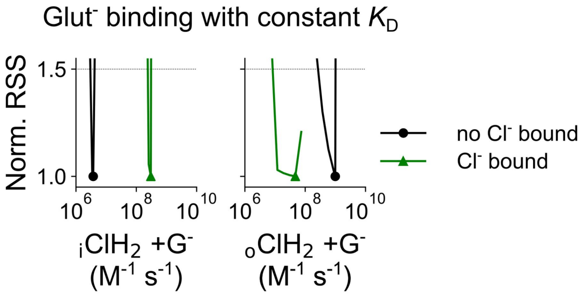


**Figure 12—figure supplement 2: Statistical analysis of glutamate binding rates, assuming a constant $K_D$. A,** Changes in the goodness of fit upon modification of the binding rate for glutamate with and without $Cl^-$, in the inward- and outward-facing conformation. During modification, the unbinding constants were simultaneously altered to keep the ratio (i.e. glutamate/aspartate-binding affinity) constant. Amplitudes are the optimized value plus 50 logarithmically distributed points between 1 and the ligand binding limit of $5\times10^9$; binding rates are at -160 mV and normalized to 140 mM glutamate or aspartate. The same RSS was derived from a wide range of rate values, indicating that the $K_D$ determines the RSS and that individual binding/unbinding rate constants do not play a major role.

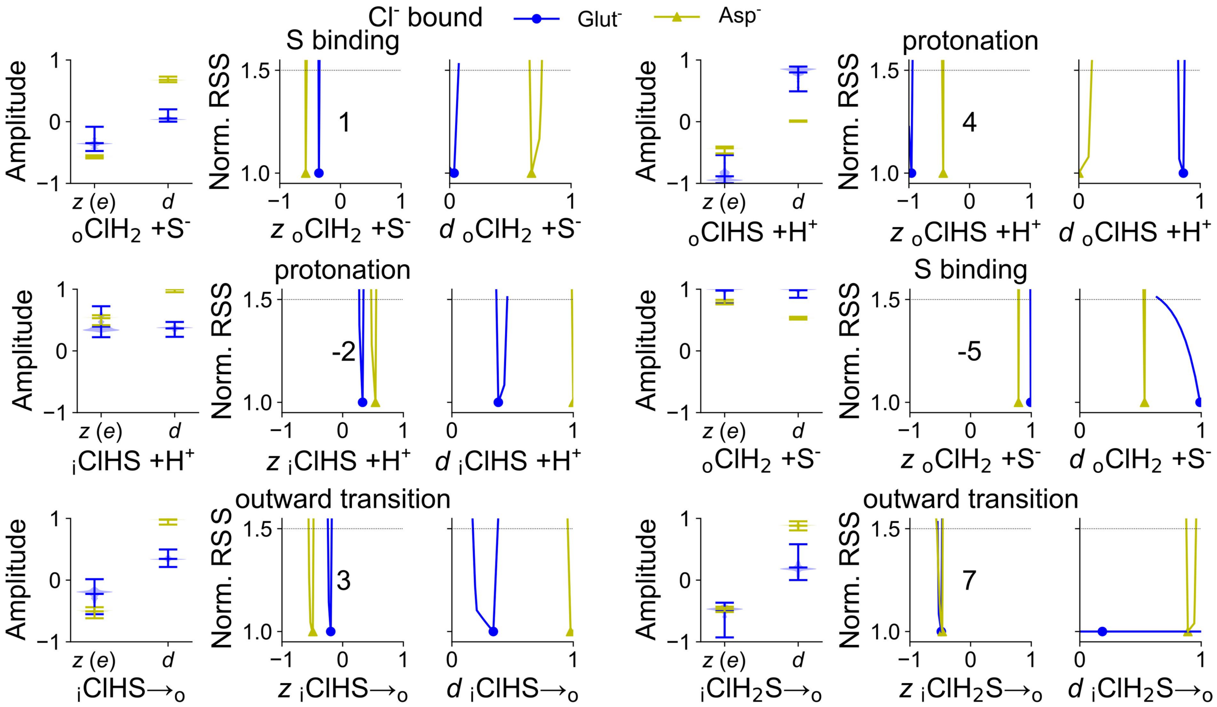


**Figure 13—figure supplement 1: Voltage dependence of protonation and substrate binding in the Cl$^-$-bound active transport cycles.** Distribution of *z* and *d* parameters for substrate binding (1 and 5), protonation steps (2 and 4), and transition between inward- and outward-facing conformations (3 and 7). Simulation results are given as violin plots depicting the amplitude range generated by exploratory mutation and normalized RSS representing goodness of fit for a range of amplitudes. Data for the outward transition of transport cycle step 6 (Figure 13) was omitted due to being substrate-independent.

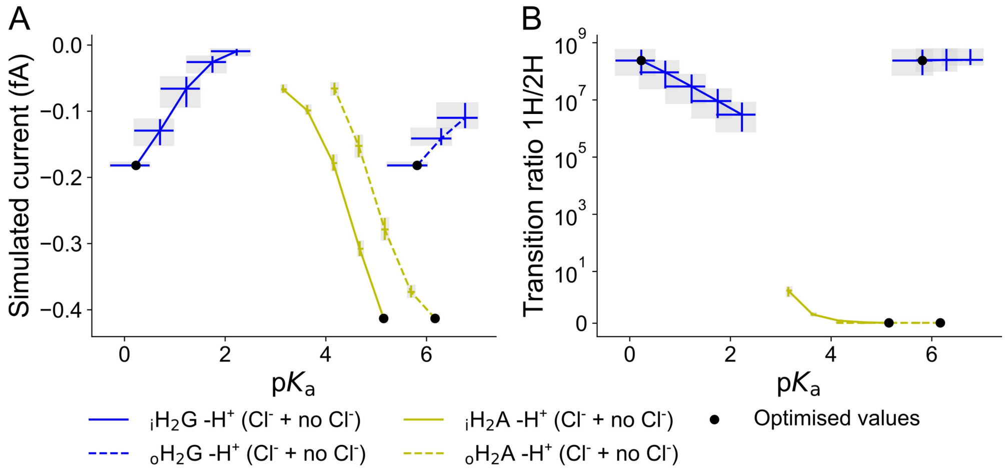


**Figure 13—figure supplement 2: Ligand-bound deprotonation rates are the major determinants of transport rates.** Predicted glutamate (blue) and aspartate (yellow) unitary currents (given as the number of charges per second in steady state × elementary charge; **A**) or relative number of $H^+$-substrate exchange transport cycles (**B**) upon modification of the optimized $pK_a$ value for the second protonation in the inward- or outward-facing substrate-bound conformation by increasing or decreasing the deprotonation rate constants by a factor of 3, 10, 33, or 100 while maintaining microscopic reversibility. Whereas the transport rates strongly depend on the $pK_a$ for both substrates, glutamate is transported in an exchange mode and aspartate in a uniport mode for all tested $pK_a$ values.

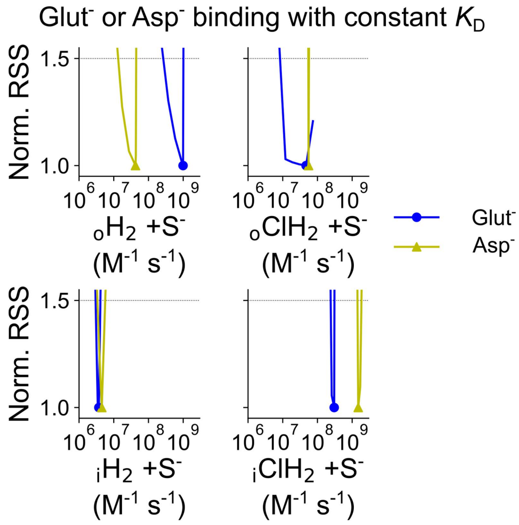


**Figure 13—figure supplement 3: Statistical analysis for substrate binding rates, assuming constant $K_D$.** Statistical analysis for substrate binding rates, assuming constant $K_D$. Changes in the goodness of fit upon modification of the binding rate for glutamate and aspartate, with and without $Cl^-$, in both conformations. During modification unbinding constants were simultaneously altered to keep the ratio; i.e. the glutamate/aspartate binding affinity, unaltered. Amplitudes are the optimized value plus 50 logarithmically distributed points between 1 and the ligand binding limit of $5\times10^9$, binding rates are at -160 mV and normalized to 140 mM glutamate or aspartate. The same RSS caused by a wide range of rate values indicates the $K_D$ determines the RSS, while individual binding/unbinding rate constants play no major role.

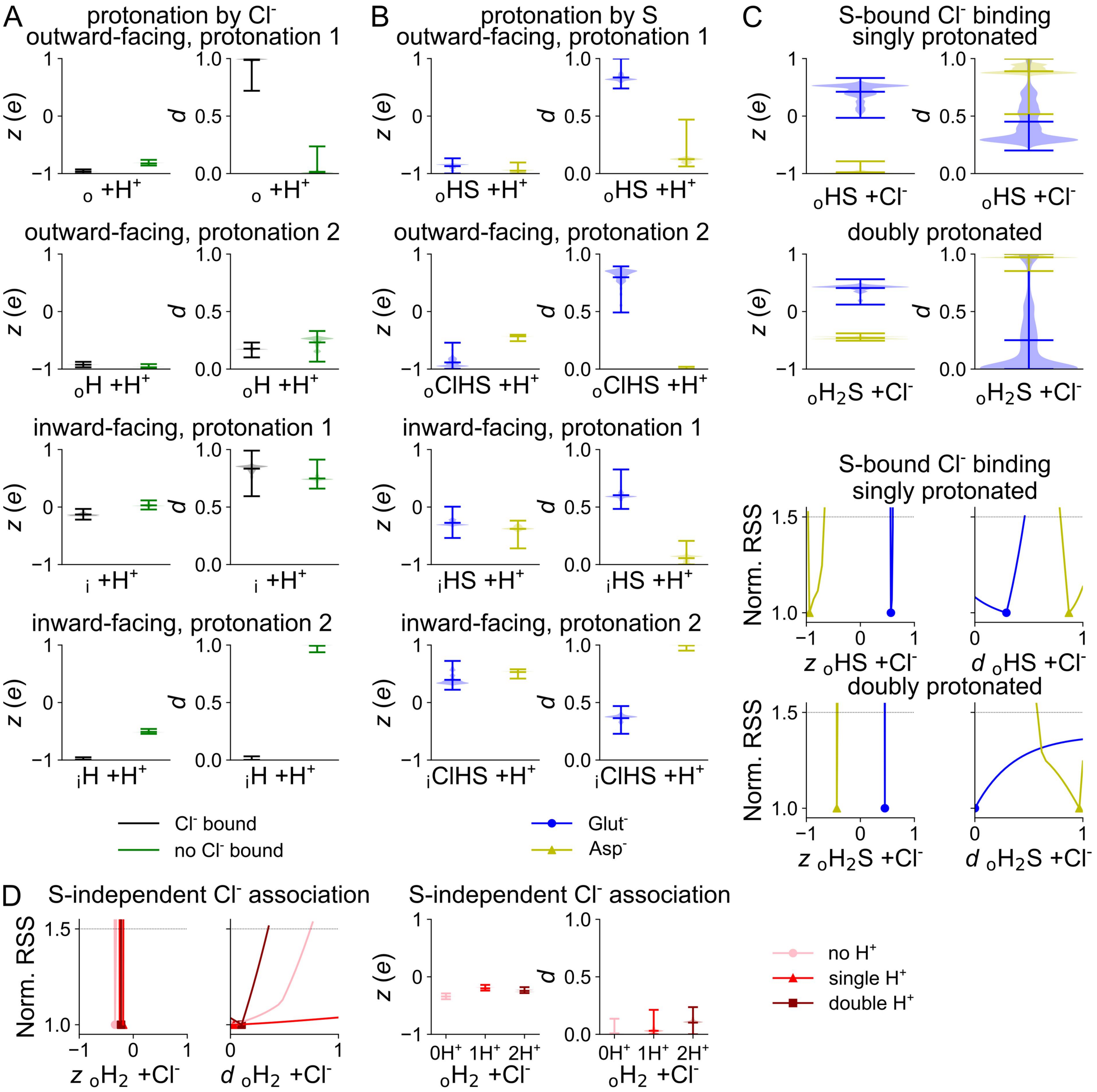


**Figure 14—figure supplement 1: Voltage dependence of protonation and $Cl^-$ binding in the $VGLUT1_{PM}$ active transport model.** Parameters for the first and second protonation without bound glutamate or aspartate for inward- and outward-facing conformations, as modulated by external $Cl^-$ (**A**), first and second protonation with substrate bound for inward- and outward-facing conformations (**B**), $Cl^-$ binding with substrate bound and with single or double protonation (**C**), and $Cl^-$ binding with no substrate bound and with no, single, or double protonation (**D**). Protonation parameters are represented by violin plots, other simulation results are given as normalized RSS representing goodness of fit for a range of amplitudes in addition to violin plots depicting the amplitude range generated by exploratory mutation.

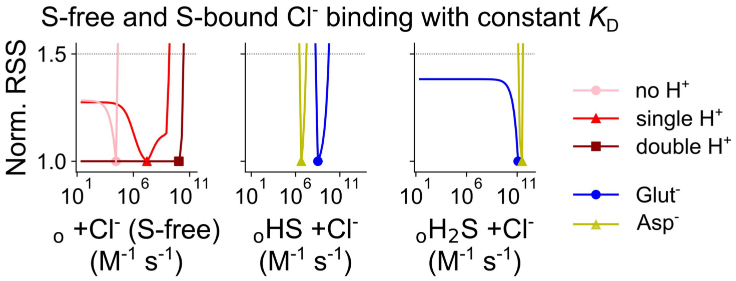


**Figure 14—figure supplement 2: Statistical analysis for $Cl^-$-binding rates, assuming a constant $K_D$.** Changes in the goodness of fit upon modification of the $Cl^-$-binding rates with no substrate bound, by protonation state in shades of red (left), or with glutamate or aspartate (substrate) bound for the single and double protonation states (middle, right). During modification, unbinding constants were simultaneously altered to keep the ratio (i.e. the binding affinity) constant. Amplitudes are the optimized value plus 50 logarithmically distributed points between 1 and the ligand binding limit of $5\times10^9$; binding rates are at -160 mV and normalized to a [$Cl^-$] of 40 mM. The same RSS derived from a wide range of rate values indicates that the $K_D$ determines the RSS, with no major role for individual binding/unbinding rate constants.
